# The Human Milk Oligosaccharide 3’Sialyllactose Promotes Inflammation Resolution and Reduces Atherosclerosis Development in Mice

**DOI:** 10.1101/2021.03.19.433472

**Authors:** Ariane R. Pessentheiner, Nathanael J. Spann, Chloe A. Autran, Bastian Ramms, Austin W.T. Chiang, Kaare V. Grunddal, Yiwen Wang, Anthony Quach, Laela M. Booshehri, Alexandra Hammond, Chiara Tognaccini, Joanna Latasiewicz, Joseph L. Witztum, Hal M. Hoffman, Nathan E. Lewis, Christopher K. Glass, Lars Bode, Philip L.S.M. Gordts

## Abstract

Macrophages contribute to the induction and resolution of inflammation and play a central role in the chronic low-grade inflammation in cardiovascular diseases caused by atherosclerosis. Human milk oligosaccharides (HMOs) are complex unconjugated glycans unique to human milk that benefit infant health and act as innate immune modulators. Here, we identify the HMO 3’sialyllactose (3’SL) as a natural inhibitor of TLR4-induced low-grade inflammation in macrophages and endothelium. Transcriptome analysis in macrophages revealed that 3’SL attenuates a selected set of inflammatory gene expression and promotes activity of LXR and SREBP. These acute anti-inflammatory effects of 3’SL were associated with reduced histone H3K27 acetylation at a subset of LPS-inducible enhancers distinguished by preferential enrichment for CTCF, IRF2, BCL6, and other transcription factor recognition motifs. In a murine atherosclerosis model, both subcutaneous and oral administration of 3’SL significantly reduced atherosclerosis development and the associated inflammation. This study provides evidence that 3’SL attenuates inflammation by a transcriptional mechanism to reduce atherosclerosis development in the context of cardiovascular disease.

## INTRODUCTION

Cardiovascular diseases (CVD) are globally the leading cause of death for both men and women. The majority of CVD-related deaths are caused by complications of atherosclerosis, a complex chronic inflammatory disorder mediated by both adaptive and innate immunity. Atherosclerosis initiates when circulating low-density lipoproteins (LDL) are trapped in the subendothelial extracellular matrix of arteries where they get modified into oxidized LDL (oxLDL). oxLDL activates the overlying endothelium, promoting infiltration of monocytes that differentiate into macrophages which internalize the oxLDL and turn into cholesterol-loaded foam cells. The retained foam cells promote a low-grade production of cytokines and chemokines that directs infiltrating monocytes to distinct phenotypic programs designated as activated pro-inflammatory macrophages or resolving macrophages^1^. The transition to a pro-inflammatory phenotype is favored by hypoxia^2^, lipopolysaccharides (LPS), cholesterol crystals, as well as cytokines, such as interferon (IFN)-γ and tumor necrosis factor (TNF) together inducing toll-like receptor 4 (TLR4) signaling and activation of the inflammasome (NLRP3)^3–6^. This amplifies a chronic inflammatory response and induces pro-inflammatory cytokine secretion such as interleukin (IL)-1β, IL-6 and TNF as well as resolving cytokines among others^3, 7, 8^.

Given that low-grade inflammation plays a key role in all stages of atherosclerosis, preventing further infiltration and production of pro-inflammatory macrophages and their subsequent cytokine secretion are considered as highly promising therapeutic strategies^9–12^. Many preclinical studies provided evidence that lowering systemic inflammation or promoting inflammation resolution would promote prevention of major adverse cardiovascular events (MACE)^13–15^. As an example, genetic inactivation of *Il-1β* and *Tnf* in Western diet-fed Apolipoprotein (Apo)E-and LDL receptor (*Ldlr*)-deficient mice reduced atherosclerosis development^16–19^ as did, administration of an IL-1β neutralizing antibody^19^. Conversely, exogenous administration of IL-1β in pigs and IL-6 in mouse models increased the atherosclerosis burden^20, 21^. However, studies in *Apoe^-/-^* mice have observed that inactivation of IL-1β can worsen plaque stability and outward plaque remodeling^22, 23^. The recently completed CANTOS trial was the first large-scale proof-of-concept trial testing quarterly administered canakinumab, a neutralizing IL-1β monoclonal antibody, to a very high-risk cohort of post-myocardial infarction patients with reduced LDL-cholesterol and elevated high-sensitivity C-reactive protein (hsCRP) levels. Treated patients experienced a 15% reduction in MACE, without affecting all-cause and CVD mortality. A subsequent CANTOS analysis showed a 26-31% reduction in CVD and all-cause mortality in the subset of subjects on canakinumab who achieved a greater than average on-treatment reduction in cytokine levels (hsCRP and IL-6)^11, 12^. The studies imply that the clinical efficacy of canakinumab relies on the magnitude of the achieved cytokine reduction. Despite the promising results, the adverse effects, very high costs and administration via injections warrant searches for additional novel, safe and effective therapeutics targeting chronic inflammation that will benefit a greater CVD patient population.

Human milk oligosaccharides (HMOs) are a natural and abundant component of human milk that have a variety of biological functions shown to promote development and regulate immune function (Bode 2012; Brugman et al. 2015; Charbonneau et al. 2016). All HMOs are unconjugated glycans that carry lactose at their reducing end. Lactose can either be fucosylated to yield fucosyllactoses, sialylated to yield sialyllactoses, or elongated and branched to yield a total of more than 150 distinct oligosaccharides, each with potentially different structure-dependent activity profiles. Unlike most oligosaccharides, such as lactose, HMOs resist the low pH in the stomach as well as digestion by pancreatic and brush border enzymes^24–26^. Approximately 1% of ingested HMOs amount is absorbed, reaches the systemic circulation^27, 28^ and is excreted in the urine^29–31^. In addition, HMOs are also present in the circulation of pregnant and lactating women^32^ as well as in amniotic fluid^33^. Originally, HMOs were considered prebiotics that help shape the gut microbiome with health benefits for the breastfed infant. It has become increasingly clear that HMOs also have beneficial immunomodulatory properties independent of the gut microbiome^34, 35^. HMOs act locally on cells of the mucosa-associated lymphoid tissues and absorbed HMOs can act on a systemic level. Most reports attribute anti-inflammatory properties to HMOs, including reduction of IL-4 and increased IL-10 production in lymphocytes from adults with peanut allergy^36^, reduction in leukocyte rolling and adhesion on TNF-activated endothelium^37^, reduction in platelet-neutrophil complex formation and neutrophil activation^38^, promotion of antigen-presenting cell activation to produce anti-inflammatory chemokines and cytokines^39^ and reducing inflammatory response in an experimental arthritis model^40, 41^. However, the effects of HMOs and their mechanism of action remain uncharacterized in the majority of low-grade chronic inflammatory diseases in adults such as atherosclerosis.

In this study, we tested the hypothesis that a specific HMO attenuates atherosclerosis low-grade macrophage inflammation and atherosclerosis development in mice. Using an unbiased screen, we identified one specific HMO called 3’sialyllactose (3’SL) that was most effective in reducing macrophage inflammation while other, structurally distinct HMOs had no effect. 3’SL significantly reduced IL1β and IL6 expression in LPS-activated macrophages, in both cell lines and primary cells of both murine and human origin. Mechanistically, we provide evidence that 3’SL also accelerates the resolution phase of inflammation by influencing the recruitment of LXR/SREBP to enhancers of enzymes involved in production of resolving lipids. Importantly, our studies show that 3’SL administration reduced the development of atherosclerotic lesions in a preclinical murine model.

## RESULTS

### Human Milk Oligosaccharides Attenuate Low-Grade Inflammation

Several studies support immunomodulatory properties of HMOs in different disease models. We determined the potential of HMOs to attenuate low-grade inflammation relevant for atherosclerosis in the RAW 264.7 murine macrophage cell line. To test this, we measured IL-1β and IL-6 production in RAW 264.7 macrophages stimulated with or without a low-grade dose (10 ng/mL) of the TLR4 activator LPS^42, 43^. We co-treated the RAW 264.7 macrophages with PBS or a low (100 µg/mL) or high (500 µg/mL) dose of LPS-free HMOs pooled from different donors (pHMOs) to capture the entire variety and chemical space of HMOs that vary between individual women (Fig. 1a). Co-incubation of LPS with pHMO significantly reduced expression of pro-inflammatory cytokines *Il-1β, Il-6, Il-10* and *Tnf* in a dose depended manner compared to LPS control (Fig. 1a-b). The most potent inhibition was observed at the higher pHMO concentration (500 µg/ml) resulting in ∼70% and ∼80% reduction in LPS-induced *Il-1β* and *Il-6* mRNA levels, respectively, compared to LPS alone. This translated in a ∼80% reduction in IL-6 protein secretion after 6- and 24-hours (Fig. 1c). The results confirm that pooled HMOs can attenuate low-grade inflammation in macrophages.

**Figure 1:**
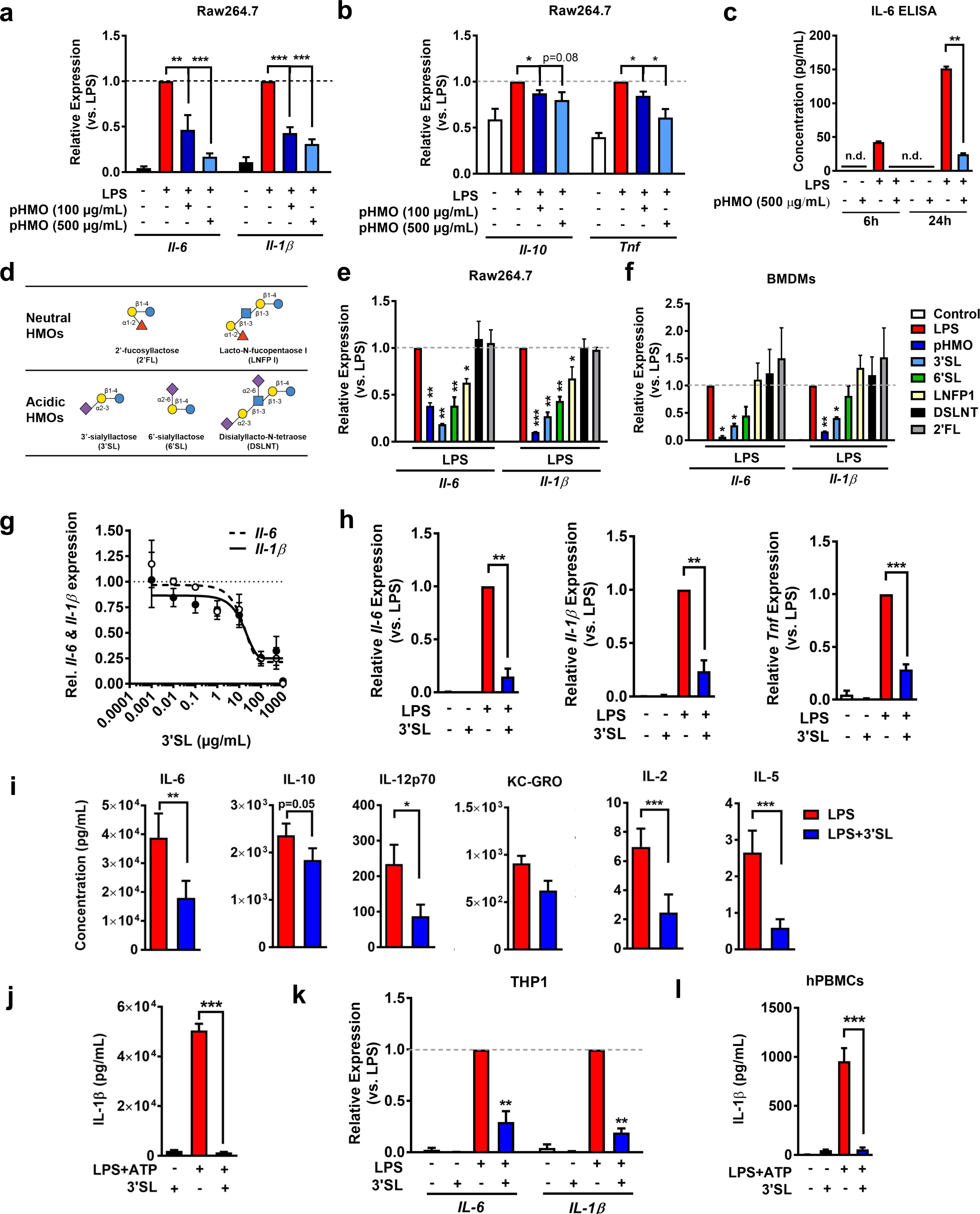
HMOs, particularly 3’sialyllactose (3’SL), reduce inflammatory cytokine expression in LPS-activated murine macrophages and human monocytes. **a-b**, Relative expression of *Il-6* and *Il-1β* (a), and *Il-10* and *Tnf* (b) in Raw264.7 cells with LPS ± pooled HMO (pHMO) incubation (n = 2). c, IL-6 protein release 6 and 24h after LPS ± pHMO incubation (n = 3). d, Depiction of individually used HMOs. e-f, Relative expression of *Il-6* and *Il-1β* in Raw264.7 (e) and murine bone marrow derived macrophages (BMDMs) (f) when treated with individual HMOs (n = 2). g, Dose response curve of 3’SL in BMDMs (n = 3). h, Relative expression of *Il-6, Il-1β,* and *Tnf* expression in PBS, 3’SL or LPS ± 3’SL (100 µg/mL) treatment. (n = 3-4). i, Cytokine concentrations in the conditioned medium of BMDMs with 24 hours LPS ± 3’SL incubation (n = 4). j, Relative expression of *IL-6* and *IL-1β* in LPS-activated human THP-1 cells treated ± 3’SL (n = 3). k, IL-1β protein release 24 hour after LPS+ATP co-incubation ± 3’SL (n ≥ 5). l, IL-1β protein release 24 hour after LPS+ATP co-incubation ± 3’SL (100 µg/mL) in human peripheral blood monocytes (hPBMCs) from three healthy donors. One-way ANOVA or unpaired student *t*-tests were used to calculate statistical significance between treatments. If not otherwise stated incubations lasted for 6 hours with 10 ng/mL LPS and 100 µg/mL 3’SL or above indicated concentrations of HMOs. (* p<0.05; ** p<0.01; *** p<0.001); Data represented as mean ± SEM.

### 3’Sialyllactose Inhibits Inflammation in Human and Murine Macrophages

Next, we attempted to identify the most potent specific HMO responsible for attenuating LPS-induced inflammation in RAW 264.7 macrophages. First, we applied anion exchange chromatography to fractionate pHMO by charge into neutral (non-sialylated) and acidic (sialylated) HMOs. RAW 264.7 macrophages were co-treated with LPS (10 ng/ml) and either neutral HMOs, acidic HMOs or lactose (each 500 µg/ml) (Extended Data Fig. 1a-b). Lactose did not affect inflammatory cytokine expression in RAW 264.7 macrophages (Extended Data Fig. 1a-b). Both predominantly acidic and neutral pHMO fractions were able to significantly decrease LPS-induced expression of *Il-1β* and *Il-6* (Fig. Extended Data 1a-b). However, crude neutral HMOs contained a small amount of the acidic HMO, 3’SL, and acidic HMOs contained small amounts of lactose and the two neutral HMOs 2’fucosyllactose (2’FL) and difucosyllacto-*N*-hexaose (DFLNH) (Extended Data Fig. 1b-c). Therefore, we tested LPS-free pHMO (500 µg/ml) alongside individual HMOs from both subfractions, namely 2’FL, difucosyllacto-*N*-tetraose (DFLNT), lacto-*N*-fucopentaose I (LNFP I), 3’sialyllactose (3’SL), and 6’sialyllactose (6’SL) each at a concentration of 100 µg/ml (Fig. 1d). The two sialylated HMOs 3’SL and 6’SL had the strongest *Il-6* and *Il-1β* inhibition in LPS-activated macrophages, while fucosylated 2’FL did not exhibit anti-inflammatory properties (Fig. 1e). Importantly, 3’SL-mediated anti-inflammatory effects observed in RAW 264.7 macrophages were confirmed in murine primary bone marrow derived macrophages (BMDMs) (Fig. 1f). Following these results, we identified 3’SL as the most potent HMO. Dose-range finding studies in BMDMs identified IC50 values for 3’SL around 15 µg/mL (Fig. 1g), which is comparable to 3’SL plasma concentrations in breastfed infants^27, 28^.

Importantly, in the absence of LPS, incubation with 3’SL (100 µg/ml) did not affect cytokine expression compared to PBS (Fig. 1h). The inhibition of IL-6, IL-10, IL-12p70, IL-2, IL-5, and IL-1β was confirmed on the protein level in culture medium of LPS-activated BMDMs co-treated with 3’SL for 24 hours, while KC-GRO was unaffected (Fig. 1i-j). Importantly, the anti-inflammatory effects of 3’SL translated to human immune cells as 3’SL also attenuated *IL-6* and *IL-1β* mRNA expression in LPS-stimulated THP1 human monocytic cells (Fig. 1k), as well as protein secretion of IL-1β in human peripheral blood monocytes (hPBMCs) (Fig. 1l). Taken together, we identified that the HMO 3’SL effectively reduced low-grade inflammatory cytokine production in murine and human macrophages and monocytes stimulated with LPS.

### 3’Sialyllactose Does not Alter TLR4 Activation and NF-κB Signaling

To gain insight into the potential mechanism involved in 3’SL-mediated reduction of pro-inflammatory cytokines, we explored several possible cell surface interactors (Fig. 2a). First, we tested whether 3’SL directly inhibits LPS binding to the TLR4 receptor through interaction with the carbohydrate moiety of LPS^42, 43^. Therefore, we activated BMDMs with the lipid portion of LPS (lipid A). Activation of *Il-6* and *Il-1β* expression was comparable to LPS, and 3’SL attenuated the expression of both cytokines in the presence of lipid A to the same extend as LPS by ∼90% and ∼70%, respectively (Fig. 2b). The results suggest that 3’SL attenuates LPS-driven inflammation without inhibiting carbohydrate interactions. Further Western blot analysis revealed that the time-dependent phosphorylation of the key signaling molecules NF-κB (p-p65) and mitogen-activated protein kinases (MAPK, p-p38) pathways were not altered by 3’SL co-incubation with LPS (Fig. 2c), suggesting that 3’SL does not directly impact the TLR4-mediated signaling cascade.

**Figure 2:**
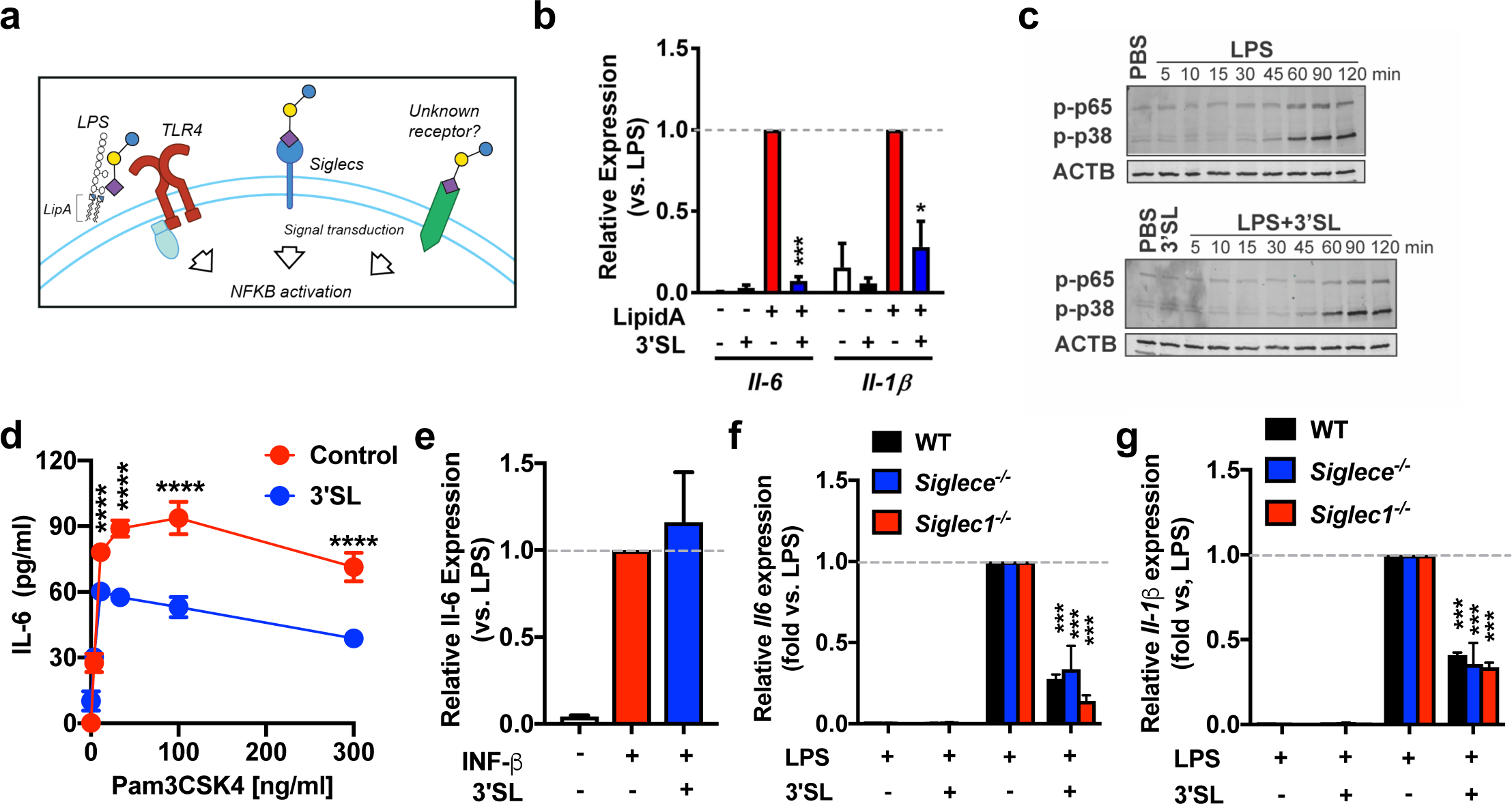
3’SL does not engage with carbohydrate part of LPS, neither reduce NF-kB signaling, nor INF-y signaling and Siglec interactions. **a**, Scheme of potential 3’SL effector pathways. Toll-like receptor 4, (TLR4) b, Relative *Il-6* and *Il-1β* mRNA expression in lipid A activated BMDMs treated ± 3’SL. c, Western blot of phospho-p65 and p38 after different times of LPS ± 3’SL stimulation. One representative blot of n = 3. d, IL-6 concentrations in the conditioned medium of BMDMs with 6 hours Pam3CSK4 (at indicated doses) ± 3’SL incubation (n = 4). e, Relative *Il-6* and *Il-1β* mRNA expression in BMDMs stimulated with 10 ng/mL interferon beta INF-β ± 3’SL. f-g, Relative *Il-6* and *Il-1β* mRNA expression in *Siglec1^-/-^* (f) and *SiglecE^-/-^* (g) BMDMs stimulated with LPS ± 3’SL. All stimulations 10 ng/mL LPS, 100 µg/mL 3’SL (* p<0.05; ** p<0.01; *** p<0.001, **** p<0.0001); Bar plots represent mean ± SEM; (n = 2-3) of individually isolated BMDMs.

Furthermore, IL-6 production in BMDMs after stimulation with the TLR2 agonist, Pam3CSK4, was also significantly attenuated 1.5-fold by 3’SL co-incubation (Fig. 2d). Upon LPS-activation in macrophages, the TLR4/MD-2 complex becomes internalized into endosomes and triggers signaling cascades that activate the expression of type I interferons (IFN) such as IFN-β^44^. IFNs are critical for innate immune responses^44, 45^ and implicated in the pathogenesis of chronic inflammatory diseases such as atherosclerosis^8, 46, 47^. Therefore, we tested if 3’SL also alleviates IFN-mediated inflammation. Exogenous stimulation of BMDMs with *IFN-β* induced *Il-6*, but not *Il-1β* mRNA expression (Fig. 2e). Co-incubation of *IFN-β* with 3’SL did not attenuate *Il-6* expression, indicating that anti-inflammatory effects of 3’SL are not mediated by attenuating the down-stream *IFN-β*-dependent pathway induced by TLR4 activation. Thus, 3’SL attenuates inflammatory gene expression in TLR4 activated macrophages without affecting TLR4 induction and NF-κB signaling.

### Anti-inflammatory Properties of 3’SL are Independent of Sialic Acid Binding Siglec Receptors

There are two main sialic acid-binding transmembrane receptors presented at the cell surface of murine macrophages, sialic acid-binding immunoglobulin-type lectin (Siglec)-1 and Siglec-E. These receptors in particular bind terminal sialic acids to modulate inflammatory signaling via intracellular tyrosine-based signaling motifs, especially immunoreceptor tyrosine-based inhibitory motifs (ITIMs) that are implicated in cell signaling and endocytosis^48, 49^. Siglec-E preferentially interacts with sialic acid linked in the *α*2-3 position to D-galactose, such as 3’SL. Sialoadhesin (Siglec-1) shares the substrate-specificity with Siglec-E and has been shown to also bind 3’SL^48, 50^. As 3’SL features a sialic acid at the non-reducing end of the lactose backbone, we tested these known sialic acid binding receptors as potential target receptors on macrophages that mediate the anti-inflammatory actions of 3’SL (Fig. 2f-g). We initially probed the importance of Siglec-E as Siglec-1 lacks the ITIM required to attenuate inflammation^6^. BMDMs derived from *Siglece*-deficient (*Siglece^-/-^*) mice were stimulated with PBS or 3’SL in the presence or absence of LPS.

Similar as in wildtype BMDMs, 3’SL significantly reduced LPS-induced *Il*-6 and *Il-1β* expression by 74% and 65%, respectively (Fig. 2f-g). Similar results were seen using BMDMs derived from Sialoadhesin knockout mice (*Siglec-1^-/-^*), providing evidence against a direct involvement of Siglec-E and Siglec-1 in the anti-inflammatory effects of 3’SL (Fig. 2f-g). Taken together, the results suggest that 3’SL does not evoke its action via carbohydrate interactions with LPS, altering the TLR4 signaling cascade, or binding to the anti-inflammatory Siglec receptors.

### 3’SL Represses and Induces a Selected Group of LPS-Responsive Inflammatory Genes

To investigate 3’SL-induced changes in macrophage gene expression throughout the course of an inflammatory response, we subjected BMDMs treated for 6 hours with or without (10 ng/ml) LPS in the presence or absence of (100 µg/ml) 3’SL to whole transcriptome (RNA-Seq) analysis. RNA-Seq results showed that using a 1.5-fold difference and false discovery rate of <0.05, 3’SL altered the expression of 133 genes under LPS stimulation and 53 genes under basal conditions (Fig. 3a-b). In contrast, LPS activation induced changes in over 5,000 genes using these same criteria (Extended Data Fig. 2a). The subset of the 79 genes downregulated by 3’SL under LPS stimulation are categorized by gene ontology analysis in the NF-κB Inflammatory response (Fig. 3a,c). We define this subset of 3’SL-regulated genes as 3’SL inflammatory repressed genes that include *Il-6*, *Tnf* and inflammatory markers, such as endothelin 1 (*Edn1*), chemokine C-X-C motif ligand 1-3 (*Cxcl1-3),* prostaglandin endoperoxide synthase 2 (*Ptgs2,* also known as *Cox2*) and the inflammatory mediator serum amyloid A3 (Saa3) (Fig. 3c)^51–53^. The exonic distribution of normalized tag counts for representative genes is illustrated in Fig. 3d-e and Extended Data Fig. 2d-e and qPCR further confirms attenuated gene expression of inflammatory marker genes (Extended Data Fig. 2f). The 3’SL inflammatory repressed genes do not overlap with genes downregulated in BMDMs under basal non-inflammatory conditions (Extended Data Fig. 2a-c). However, this latter 3’SL downregulated basal gene set also includes inflammation annotated genes such as fatty acid binding protein 4 (*Fabp4*), PYD and CARD domain containing (*Pycard*),

**Figure 3:**
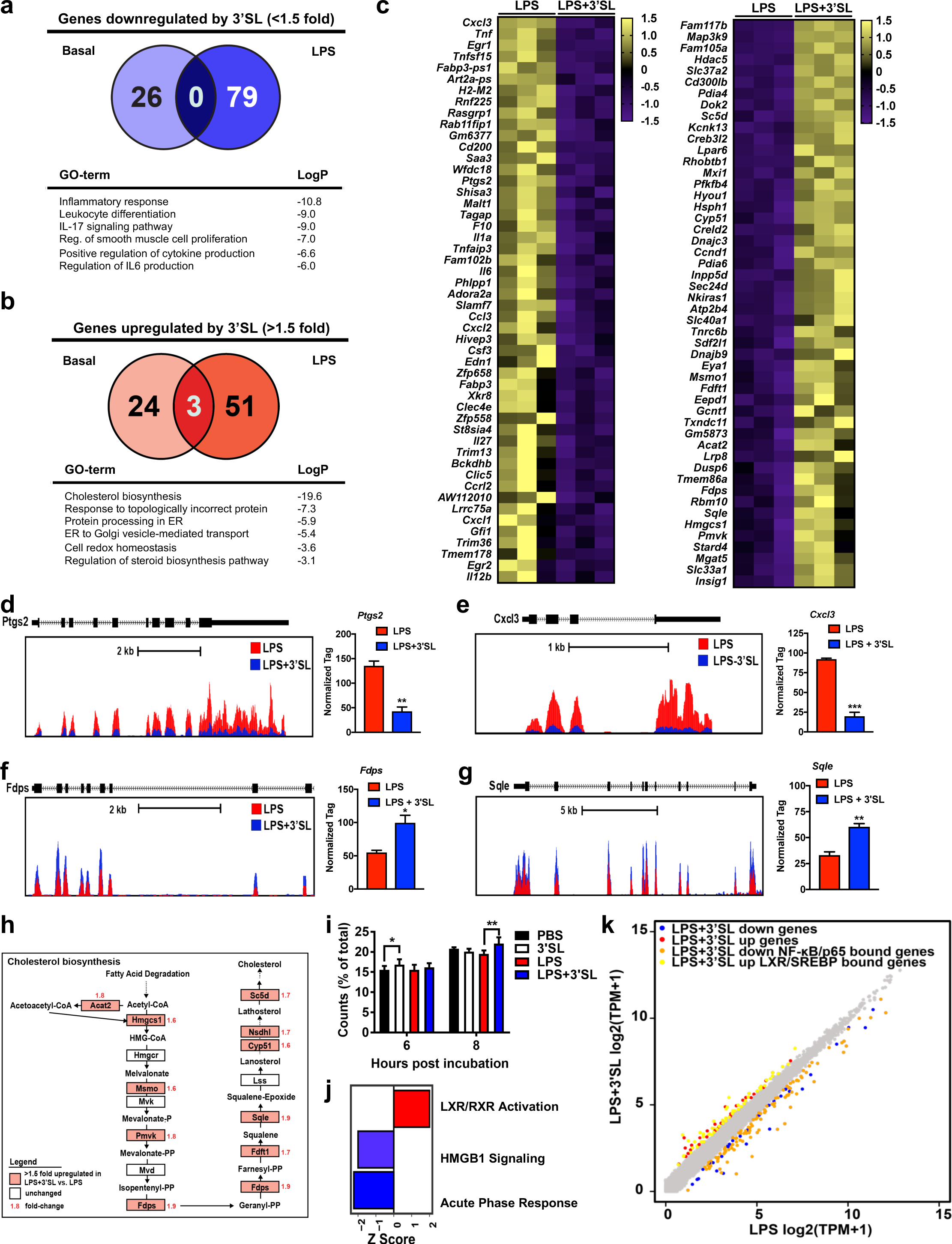
3’SL downregulates inflammatory pathways and upregulates cholesterol biosynthesis and efflux in LPS-stimulated BMDMs. **a-b**, Venn-diagrams of differentially expressed genes detected by RNA-Seq (cut-off p < 0.05 and fold change, FC, >1.5) of 3’SL down-regulated (a) and up-regulated (b) genes in LPS-activated BMDMs compared to quiescent, PBS treated BMDMs. Corresponding pathway analyses of LPS vs. LPS+3’SL treated BMDMs are below the diagrams (n = 3). c, Heat maps representing z-normalized row expression of each gene for RNA-Seq from independent biological duplicates showing the 50 most up-regulated (right) or down-regulated (left) genes in BMDMs at 6 h after LPS ± 3’SL stimulation. d-e, UCSC genome browser images illustrating normalized tag counts for *Ptgs2* and *Cxcl3* with normalized tag count averages (n = 3). f-g, UCSC genome browser images illustrating relative expression levels for cholesterol biosynthesis genes *Fdps* and *Sqle* with normalized tag count averages (n = 3). h, Scheme of KEGG-pathway cholesterol biosynthesis. Red marked genes are upregulated by 3’SL co-incubation with LPS. Red numbers above gene names indicate fold-change. i, Cholesterol efflux from stimulated BMDMs to human HDL (25 µg/mL) as an acceptor after 6 and 8 hours, respectively (n ≥ 3). j, Enriched pathways identified by Ingenuity Pathway Analysis (IPA) software. All significantly, differentially expressed genes of LPS vs. LPS+3’SL RNA-Seq data sets were used for the analysis with IPA (cut-off > 1.5-fold) with Z-score ≥ ±2. k, Scatter plot depicting the relationship between fold change of 3’SL-LPS repressed and 3’SL induced genes, overlaid with LXR and SREBP-associated accessible loci from a previously generated ATAC-seq data set^58^. (* p<0.05; ** p<0.01; *** p<0.001); Bar or dot plots represent mean ± SEM.

C1q and tumor necrosis factor related 12 (*C1qtnf12*), and Cd180 (Extended Data Fig. 2b-c). The observations suggest that 3’SL does not result in pan-attenuation of TLR4 induced changes in gene expression, but rather represses a specific subset of pro-inflammatory genes.

### Sterol and Fatty Acid Metabolism Genes are Upregulated by 3’SL During Inflammation

During LPS stimulation, 3’SL induced an almost equal number of genes compared to the downregulated group of genes. The majority of the 54 upregulated genes are involved in sterol biosynthesis, nuclear receptor activation, as well as ER-Golgi transport, and ER protein processing (Fig. 3b). This subset of 3’SL inflammatory induced genes has a minimal overlap with genes upregulated under basal conditions, indicating an inflammation-depended context wherein 3’SL affects these 3’SL inflammatory induced genes (Fig. 3b and Extended Data Fig. 2a-c). Interestingly, among the 3’SL inflammatory induced genes, a significant portion is involved in cholesterol biosynthesis, such as 3-hydroxy-3-methylglutaryl-CoA synthase 1 (*Hmgcs1*), farnesyl diphosphate synthase (*Fdps*), and squalene monooxygenase (*Sqle*), and fatty acid metabolism, such as Acyl-CoA desaturase 2 (*Scd2*) and acetyl-CoA acetyltransferase 2 (*Acat2*) (Fig. 3c-g & Extended Data Fig. 2d-f). Another 3’SL inflammatory induced gene is StAR-related lipid transfer protein 4, which is important for regulation of cholesterol homeostasis and membrane trafficking^54^ (Fig. 3h & Extended Data Fig. 2g). Low-density lipoprotein receptor-related protein 8 (*Lrp8*) was the most up-regulated 3’SL inflammatory induced gene and was also upregulated by 3’SL in quiescent BMDMs (Fig. 3c,f & Extended Data Fig. 2g-i). *Lrp8* encodes for an apolipoprotein E (apoE) receptor and promotes cholesterol efflux and lipoprotein clearance to exert its anti-atherogenic effects^55–57^. Based on the gene expression changes, we expected to see a change in cholesterol homeostasis in 3’SL treated BMDMs. Cholesterol efflux to ApoE-containing HDL as an acceptor was significantly increased in 3’SL treated BMDMs compared to control cells after 6 hours co-incubation and in 3’SL+LPS vs LPS treated BMDMs after 8 hours (Fig. 3i). These results support that 3’SL has a selective impact on gene expression changes during LPS stimulation. Importantly, the data also suggest that 3’SL engages transcription factors affecting genes involved in cholesterol and fatty acid homeostasis.

### The 3’SL Response is Associated with Activation of LXR and SREBP Signal Dependent Transcription Factors

Cholesterol and fatty acid metabolism genes are transcriptionally regulated by the nuclear Liver X Receptors (LXRs) and Sterol Regulating Element Binding Proteins (SREBPs) 1 and 2. Ingenuity Pathway Analysis software (IPA) of the RNA-Seq results revealed three metabolic pathways enriched or repressed by 3’SL (Z-score of >2 or <-2, respectively). Importantly, it independently identified Liver-X receptor (LXR)/Retinoid-X receptor (RXR) regulated pathways as the most significantly enriched (Fig. 3j). Importantly, *Lxra* (*Nr1r3* gene) and *Lxrb* (*Nr1h2* gene) expression levels were not influenced by 3’SL incubation (Extended Data Fig. 2h). We used previously generated ATAC-seq data from BMDMs^58^ to assign nearby putative regulatory enhancer regions to 3’SL regulated genes, as defined by accessible regions within 20 Kb of the gene body. To assess 3’SL-mediated modulation’s potential via LXR and SREBP, we overlaid these associated accessible loci of the 3’SL inflammation-induced genes with Chromatin-immunoprecipitation (ChIP) sequencing (ChIP-seq) data for LXR and SREBP obtained in BMDMs after stimulation with the LXR/SREBP agonist GW3965. In alignment with the IPA analysis, we observed that 67% of the 3’SL inflammatory induced genes are occupied by either LXR and SREBP (Fig. 3k), with the majority of these loci co-bound by LXR and SREBP. These data suggest a role for LXR and SREBP in the 3’SL mediated upregulation of lipid metabolism-related genes.

### LXR and SREBP Transcriptional Activity is Modulated by 3’SL

Previous studies support a role for LXR and SREBP responsive genes in the resolution phase of TLR-mediated inflammation^58^. To investigate the potential of 3’SL in modulating these resolving genes’ expression at the level of transcription, we probed the associated enhancer landscape by performing ChIP-seq for accepted markers of transcriptional activity such as histone 3 lysine 27 acetylation (H3K27ac) and p300, a histone acetyltransferase (HAT) in LPS treated BMDMs with or without 3’SL. H3K27 acetylation is deposited by HATs, such as p300, which are associated with transcriptional co-activators and is highly correlated with regulatory element activity^59^. To associate the H3K27ac signal with specific regulatory elements associated with 3’SL modulated genes, we overlapped H3K27ac with locus specific ATAC-seq peaks. More specifically, we first determined H3K27ac levels at ATAC-seq defined regions of 3’SL regulated gene loci (Fig. 4a). In line with the RNA-Seq data, we observed that activity at a limited subset of the LPS induced enhancers were upregulated by 3’SL co-incubation (Fig. 4b). *De novo* motif analysis using a GC-matched genomic background shows that these 3’SL inflammatory induced genes were enriched for myeloid specific lineage determining transcription factors (LDTF), such as AP-1, RUNX1, CEBPA and ETS-factor motifs, but these regions were also specifically enriched for downstream signal dependent transcription factors (SDTF) such as SREBP and LXR response elements (LXRE) (Fig. 4c-d). For 3’SL upregulated gene loci, we focused on transcriptionally accessible gene loci that demonstrated binding for LXR or SREBP (Fig. 4e). H3K27ac levels were robustly induced by 3’SL at those enhancer sites (Fig. 4e-h), as exemplified by genome browser tracks of 3’SL activated genes such as *Lrp8, Fdps, Stard4* and *Sdc2* (Fig. 4h & Extended Data Fig. 3a). Interestingly, 3’SL treatment alone allowed for significant induction of H3K27ac levels at those LXR/SREBP enhancers (Fig. 4e-h). The data indicate that 3’SL can blunt the LPS repression of LXR/SREBP regulated genes. This result suggests that 3’SL is conferring activation of these enhancers by modulating the activity of LXR and SREBP transcription factors. This concept is further support by 3’SL dependent recruitment of p300 at the LXR/SREBP enhancers (Fig. 4g). The LXR/SREBP bound and 3’SL induced enhancers are direct targets as demonstrated by p300, SREBP, and LXR recruitment in BMDMs to those sites upon treatment with GW3965 (Extended Data Fig. 3b). The observation further supports the idea that 3’SL is a bona fide modulator of SREBP and LXR transcriptional activity.

**Figure 4:**
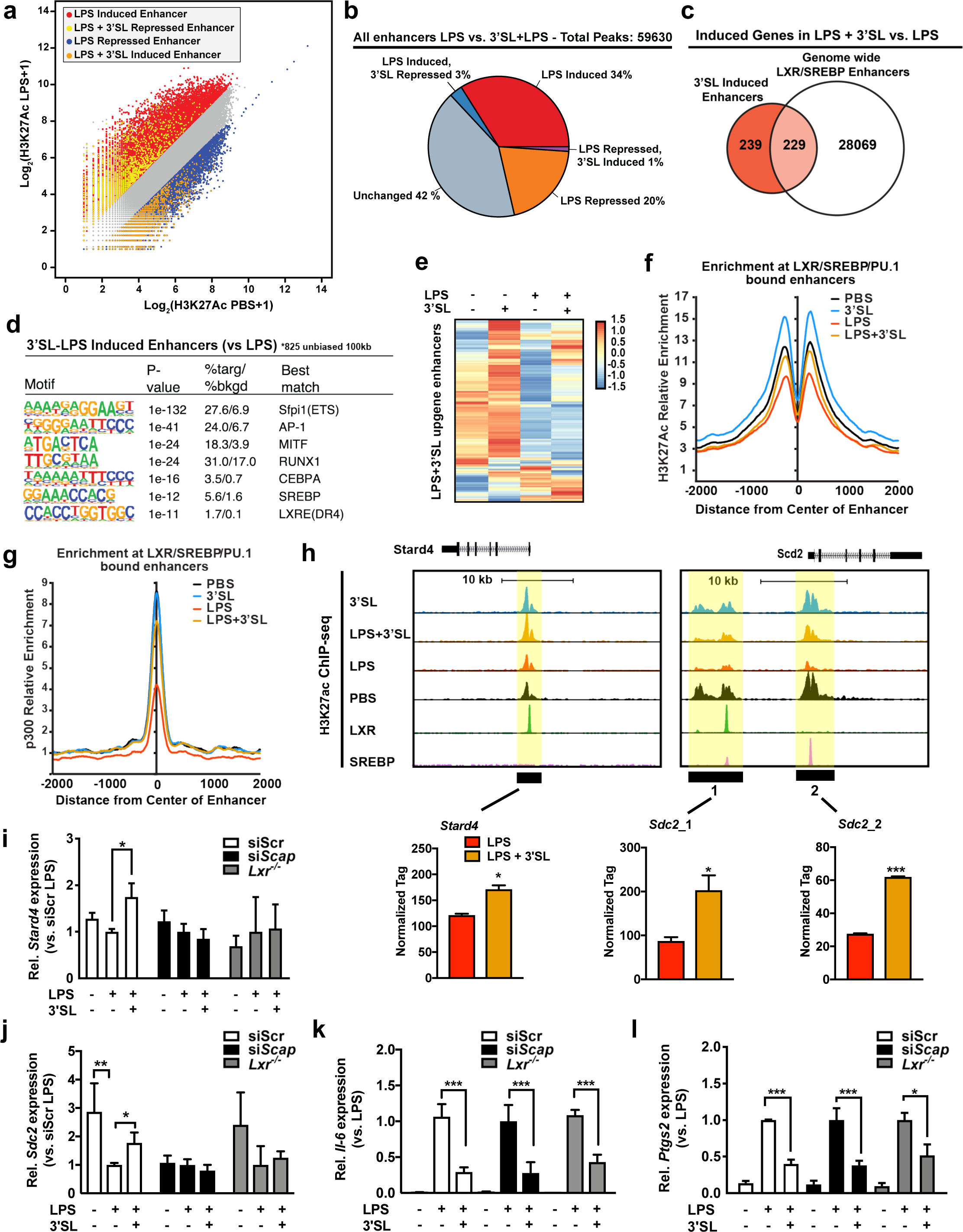
The 3’SL inflammation resolution response is associated with LXR and SREBP signal dependent transcription factors. **a**, Scatter plot depicting the enhancers as defined by H3K27ac in their relationship to LPS stimulation with and without 3’SL co-incubation. b, Venn-diagram of global enhancers affected by 3’SL and LPS compared to LPS induced enhancers. c, Venn-diagram of 3’SL-LPS induced genes compared to LPS induced enhancers associated with LXR/SREBP bound loci. d, *De novo* motif analysis of 3’SL-LPS induced enhancers (vs. LPS) using a GC-matched genomic background. e, Heatmap of the fold change in 3’SL upregulated H3K27ac levels at ATAC-seq defined gene loci that demonstrated binding for LXR or SREBP. f, Distribution of H3K27Ac tag densities, in the vicinity of genomic regions co-bound by LXR, SREBP or PU.1, in BMDMs treated with indicated stimuli for two hours. g, Distribution of p300 tag densities, in the vicinity of genomic regions co-bound by LXR, SREBP or PU.1, in BMDMs treated with indicated stimuli for two hours. h, UCSC genome browser images illustrating normalized tag counts for H3K27ac at *Stard4* and *Scd2* target loci together with mapped LXR and SREBP binding sites. i-l, Relative expression of target genes in BMDMs transfected with non-targeting control siRNA (siScr) or si*Scap* or BMDMs isolated from *Lxrα/β* knock-out mice (*Lxr*^-/-^) stimulated with PBS and LPS ± 3’SL. (n silencing experiments = 4, *Lxr*^-/-^ BMDMs from 3 individual mice). (*p<0.05; ** p<0.01; *** p<0.001); Bar or dot plots represent mean ± SEM.

Next, we set out to test if genetic inactivation of LXR or SREBP prevents the induction and repression of 3’SL inflammation modulated genes. We targeted SREBP activation by siRNA mediated knockdown of its upstream regulator SCAP1 (siSCAP; Extended Data Fig. 3f)^60, 61^. To probe the importance of LXR, we isolated BMDMs from LXRα and LXRβ double knockout mice (*Lxr^-/-^*). As expected, inactivation of LXR and SREBP blunted the induction of 3’SL inflammation-induced genes, such as *Stard4* and *Scd2* (Fig. 4i-j). In contrast, SREBP and LXR targeting did not affect the expression of 3’SL inflammation-repressed genes as 3’SL was still able to attenuate LPS-induced expression of *Il6*, *Ptgs2*, *Il1b*, and *Saa3* equally in LXR-KO and siSCAP treated BMDMs (Fig. 4k-l and Extended Data Fig. 3g). The observation supports the idea that 3’SL is a novel molecule capable of stimulating SREBP and LXR transcriptional activity in macrophages.

### 3’SL Inhibits Activation of a Subset of TLR4 Responsive Enhancers

Genes negatively affected by 3’SL belong to inflammatory pathways such as the acute phase response and the pro-inflammatory high mobility group box 1 (HMGB1) signaling (Fig. 3j) according to pathway analysis of the RNA-Seq data. We assessed the extent to which altered transcriptional regulation contributes to the 3’SL-mediated repression of inflammatory gene expression. Utilizing H3K27ac ChIP-Seq data, we evaluated the effect of 3’SL on ATAC-seq defined enhancers activated by LPS (Fig. 4a). In line with the RNA-Seq data, we observed that activity at a limited subset of the LPS induced enhancers was negatively affected by 3’SL co-incubation (Fig. 4b). *De novo* motif analysis using a GC-matched genomic background showed that these 3’SL-repressed regions demonstrate motif enrichment for the expected macrophage lineage-determining factors (PU.1, ATF3/AP-1, CEBP/NFIL3) (Fig. 5a). However, the relative order and frequencies of motifs recognized by signal-dependent transcription factors were different from those observed in the total set of LPS-induced enhancers. In particular, an ISRE recognized by IRF factors was the fourth most enriched motif in 3’SL-repressed enhancers and was present in nearly 22% of the targets. In contrast, the corresponding motif in the total set of LPS-induced enhancers ranked 5^th^ and was present in only 5% of the targets (Fig. 5a & Extended Data Fig. 4a). Conversely, motifs recognized by NF*κ*B were ranked fourth and were present in 6% of the total set of LPS-induced enhancers but ranked tenth and were present in less than 1% of 3’SL-repressed enhancers (Fig. 5a & Extended Data Fig. 4a).

**Figure 5:**
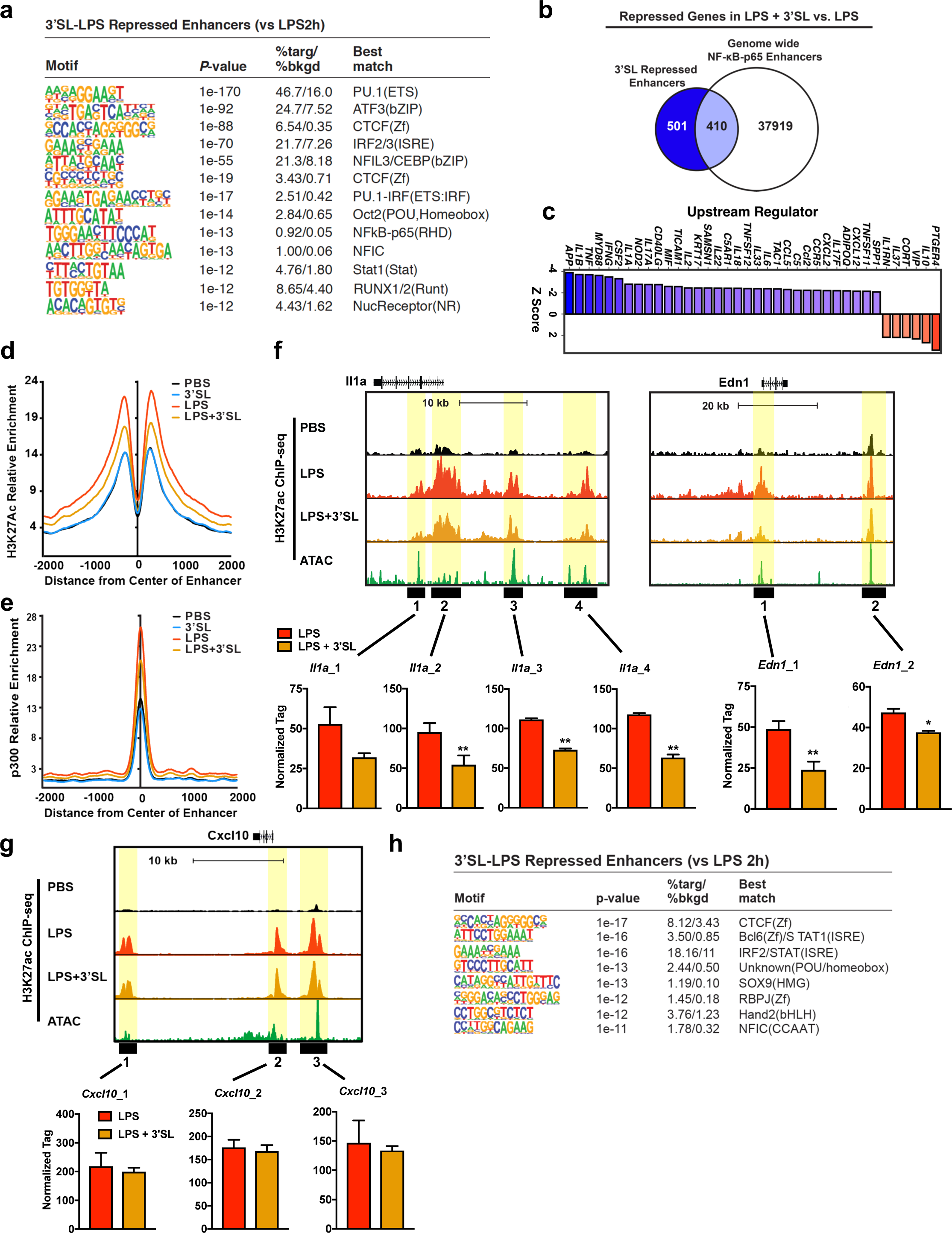
3’SL Mediates Reprogramming of the NF-κB Enhancer Landscape Activity To Attenuate Inflammation. **a**, De novo motif analysis of 3’SL+LPS repressed enhancers (vs. LPS) using a GC-matched genomic background. b, Venn-diagram of 3’SL-LPS repressed genes compared to LPS induced enhancers associated with NF-κB-p65 bound loci. c, IPA analysis of upstream regulators of differentially expressed genes (|Z-score| > ±2). d, Distribution of H3K27ac tag densities in the vicinity of genomic regions co-bound by NF-κB-p65 in BMDMs treated with indicated stimuli for two hours. e, Distribution of p300 tag densities in the vicinity of genomic regions co-bound by NF-κB-p65 in BMDMs treated with indicated stimuli for two hours. f-g, UCSC genome browser images illustrating normalized tag counts for H3K27ac *at Il1a, Edn1*, *Ifnb1 and Cxcl10,* target loci together with mapped ATAC-seq tags. h, *De novo* motif analysis of 3’SL+LPS repressed enhancers (vs. LPS) using all LPS induced ATAC-seq peaks as a background.

These observations support the concept that both the 3’SL-sensitive and insensitive enhancers are selected by a common set of macrophage lineage determining factors ^62^, but that the motifs for signal-dependent transcription factors that act upon these enhancers differ. To further explore the role of NF*κ*B, we overlapped the 3’SL-sensitive enhancers with the DNA binding pattern of the p65 subunit of NF*κ*B following LPS stimulation. This analysis indicated that despite consensus kB motifs being present at less than 1% of the 3’SL repressed enhancers, ∼44% were occupied by p65 (Fig. 5b). This result suggests that p65 binding at these locations is mediated by weak motifs and/or by indirect mechanisms. Functionality of p65 at these locations is in line with the upstream signal transduction regulatory analysis of the RNA-Seq data indicating that the pattern of inhibition mediated by 3’SL is similar to that resulting from inhibition of inducers of NF-κB responses, such as IL-1ß, IL-6, TNF, and the NF-κB activator MyD88 in LPS-stimulated BMDMs (Fig. 5c). Furthermore, LPS-dependent induction of both H3K27ac and p300 recruitment was significantly reduced at local NF-κB-p65 bound enhancers (Fig. 5d-e). 3’SL alone did not regulate H3K27ac levels or p300 recruitment at these NF-κB-p65 enhancers in the absence of LPS stimulation (Fig. 5d-e). Examples of the relationships of p65 binding to 3’SL-repressed enhancers are exemplified at the *Edn1* and *Il1a* genes in Fig. 5f. In contrast, *Cxcl10*, *Mx1*, *Socs3*, *Ifnb* represent LPS-induced genes that are not repressed by 3’SL and show no changes in H3K27ac (Fig. 5g & Extended Data Fig. 4b).

The finding that a subset of 3’SL-repressed enhancers are occupied by p65, but lack consensus kB motifs, led us to modify the motif enrichment analysis such that all LPS-induced enhancers were used as the background, rather than a GC-matched random genomic background. This approach eliminates motifs that are common to 3’SL-sensitive and 3’SL-insensitive enhancers, such as PU.1 and AP-1, and results in identification of motifs that are enriched in one subset but not the other. This analysis resulted in the identification of several motifs that occur at frequencies that are consistent with functional importance, including an ISRE (18%) recognized by IRFs, a motif recognized by the chromatin conformation modulator CTCF (8%) and a motif recognized by established repressor B cell leukemia 8 (BCL6) (4%) (Fig. 5h). Notably, a consensus NF*κ*B is not significantly enriched, supporting the concept that localization of p65 to a subset of 3’SL-sensitive enhancers is mediated by indirect interactions. Collectively, these findings provide evidence that 3’SL represses a select subset of LPS-induced genes by suppressing the activity of enhancers that lack consensus NFκB motifs and are instead regulated by distinct combinations of signal-dependent transcription factors.

### Systemic 3’SL Administration in *Ldlr^-/-^* Mice Attenuates Atherogenesis

Given the importance of chronic inflammation and inflammation resolution in atherosclerosis, we asked if 3’SL administration could be used therapeutically. We administered 3’SL via subcutaneous (s.c.) injections of Western-type diet (WTD: 42 % kcal from fat, 0.2 % total cholesterol) fed *Ldlr^-/-^* mice, as oral administration of 3’SL alters the gut microbiome and could confound interpretation of results^34, 35^. Based on the blood volume of an average mouse (77-80 ml/kg) we evaluated the pharmacodynamics of a 400 µg and 200 µg s.c. 3’SL injection in 200 µl PBS (Fig. 6a). Pharmacokinetic analysis showed that 3’SL appeared in the circulation within 5 min after s.c. injection. The concentration peaked at 20 min (20 and 35 ug/mL plasma, respectively) and returned to the baseline after 180 min (Fig. 6a). Based on the rapid clearance of 3’SL, we injected 400 µg s.c. twice daily as this concentration was within our effective IC50 dose tested *in vitro* (Fig. 1g). Male *Ldlr^-/-^* mice were given WTD two weeks prior to initiation of 3’SL treatment to induce hypercholesterolemia and this continued for the remainder of the experiment. Two-weeks after initiation of the WTD, *Ldlr^-/-^* mice were injected with 3’SL (400 µg in 200µl PBS) or PBS (200 µl) twice daily for 6 weeks (Fig. 6b). 3’SL therapy did not alter food intake, body weight gain or organ weights of adipose tissues and liver at the time of harvest compared to PBS control treated *Ldlr^-/-^* mice (Extended Data Fig. 5a-c). Plasma lipid levels and lipoprotein profiles were not altered by 3’SL intervention (Fig. 6c-d & Extended Data Fig. 5d-e). Fasting blood glucose or hepatic lipid content did not change (Extended Data Fig. 5f-g). This establishes that 3’SL administration did not alter the main atherosclerotic drivers in *Ldlr^-/-^* mice.

**Figure 6:**
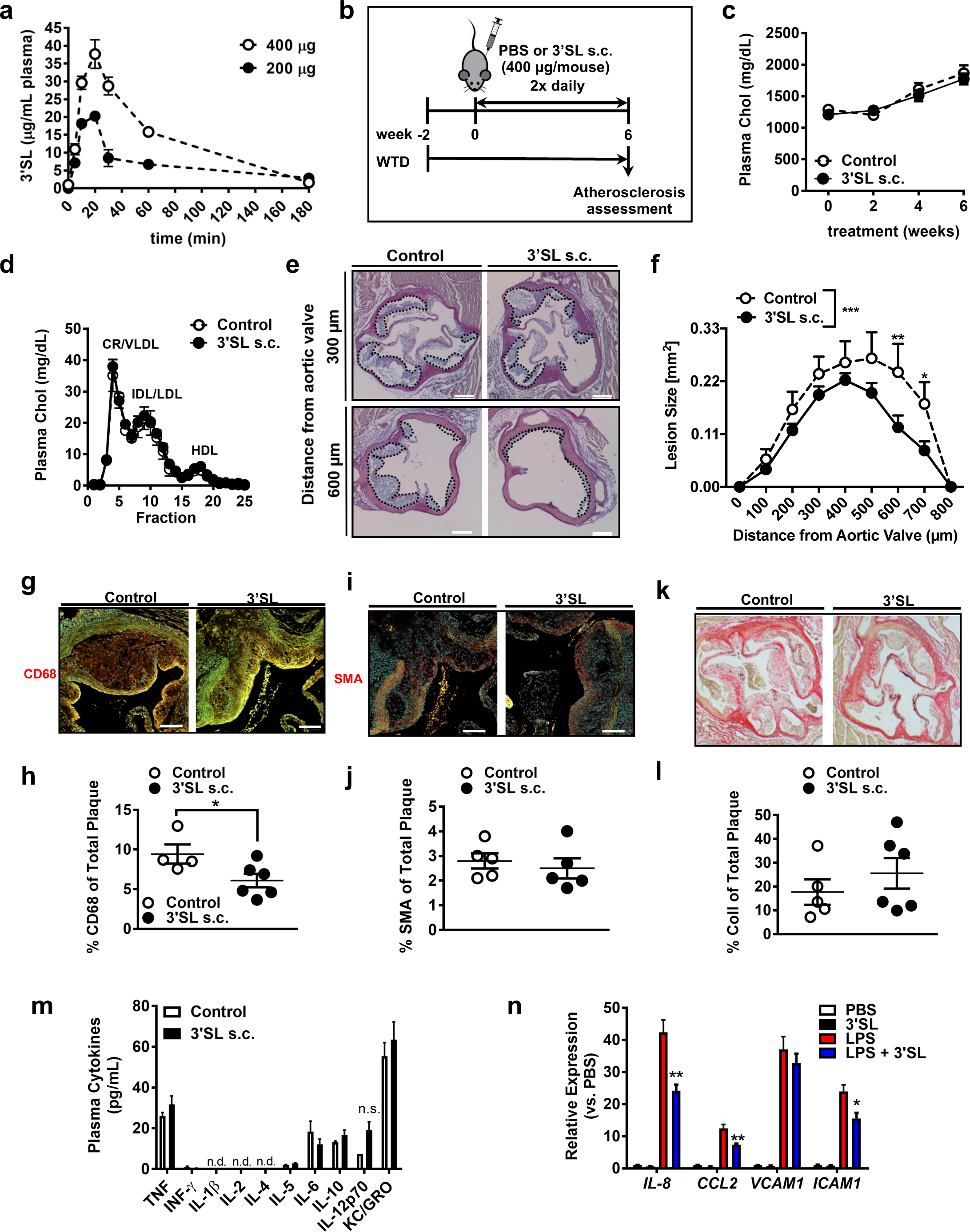
Six-week subcutaneous treatment with 3’SL leads to reduced atherosclerotic development without negative side-effects associated with body weight, plasma lipid parameters, and blood glucose. **a**, Pharmacokinetics of 3’SL after subcutaneous (s.c.) injections (n = 3 per group/time point). b, Treatment regimen. Male *Ldlr^-/-^* were put on a Western type-diet (WTD) for 8 weeks. After two weeks, mice were treated twice daily with s.c. injections of 400 µg 3’SL in PBS per mouse for 6 weeks. PBS injections served as a control. c, Bi-weekly plasma cholesterol levels in 3’SL treated and control *Ldlr^-/-^* mice (n = 14-15). d, FPLC cholesterol lipoprotein profiles after 6 weeks of treatment (two pooled samples per group of 7-8). e-f, Representative H&E staining (e) and quantification of atherosclerotic lesion size (f) in the aortic sinus (scale bar = 100 µm, n = 7-8). g-h, Atherosclerotic lesions stained with CD68 for macrophages (g) and quantification (h) of the positive stained area (n = 4-6). i-j, Smooth muscle actin (SMA) staining (i) and quantification (j) of the positive stained area (n = 5-6). k-l, Picrosirius red staining for collagen (k) and quantification (l) of the positive stained area (n = 5). m, Cytokine concentrations in the plasma after 6 weeks treatment (n = 6). n, Relative expression of inflammatory marker genes in human umbilical vein endothelial cells (HUVECs) treated with PBS or LPS (10 ng/mL) ± 3’SL (100 µg/mL) (n = 3). (* p<0.05; ** p<0.01; *** p<0.001); Bar or dot plots represent mean ± SEM.

We next examined the impact of 3’SL on atherogenesis. Atherosclerosis lesion volume was assessed via *en face* aorta and aortic root analysis (Fig. 6e-f & Extended Data Fig. 5h). The short 8-week WTD feeding regimen resulted in very little *en face* plaque in the aorta (> 2% of aortic surface area-data not shown), which was not different between treatment groups (Extended Data Fig. 5h). In contrast, analysis in the aortic root, where lesions are more advanced in mice, revealed that 3’SL treatment reduced lesion volume by 30% (1.4 ± 0.2 mm^3^ vs 0.96 ± 0.09 mm^3^; *p* = 0.002) (Fig. 6e-f). These data imply that therapeutic 3’SL administration reduces development of WTD-induced atherosclerosis in *Ldlr^-/-^* mice independent of changes in plasma lipoprotein levels.

### Intervention with 3’SL Reduces Macrophage Lesion Content and Improves Plaque Stability

We also evaluated the impact of 3’SL treatment on atherosclerotic plaque cell and extracellular matrix composition. We initially quantified lesion macrophage content via CD68 immunofluorescent staining. 3’SL therapy was associated with a significant 1.5-fold reduction in macrophage content in 3”SL mice when compared to equal-sized lesions from PBS treated *Ldlr^-/-^* mice (6.1% of lesion area versus 9.4%; p < 0.05) (Fig. 6g-h). No differences in smooth muscle cell content, and a slight trend towards increased collagen content in 3’SL treated mice were noted (Fig 6i-l). However, despite a reduction in macrophage content visual inspection of van Gieson-stained cross-sections of lesions at the aortic root did not show differences in necrotic core content between the treatment groups (Extended Data Fig. 5i). Nor did we observe alterations in systemic plasma cytokine concentrations after six weeks of treatment between the treatment groups (Fig. 6m). Plasma monocyte, neutrophil and lymphocyte counts were also unchanged (Extended Data Fig. 5j-k). Although we previously showed that 3’SL does not directly interfere with monocyte’s rolling and adhesion to activated endothelium^37^, we also evaluated if 3’SL promotes comparable anti-inflammatory properties on activated endothelium. Stimulation of human umbilical vascular endothelium (HUVEC) with (10 ng/ml) LPS induced expression of known factors that promote monocyte recruitment and invasion such as *Il-8*, vascular adhesion molecule-1 (*VCAM1*), intercellular adhesion molecule-1 (*ICAM1*) and the monocyte chemoattractant protein C-C motif chemokine ligand 2 (*CCL2*) (Fig. 6n). Treatment of HUVEC with 3’SL alone did not affect the expression of these inflammatory genes. However, co-treatment of HUVEC with LPS and 3’SL (100 µg/ml) significantly reduced the expression of adhesion molecule *ICAM1* by 1.5-fold, chemoattractant proteins *IL-8* and *CCl2* by 1.8-and 1.7-fold (Fig. 6N), respectively, supporting the idea that 3’SL could inhibit trafficking of monocytes into an inflamed artery. Overall, the findings suggest that subcutaneous 3’SL treatment attenuates atherosclerosis development by reducing the number of macrophages in the lesions.

### Oral 3’SL Administration Attenuates Atherogenesis in *Ldlr^-/-^* Mice

We next investigated if oral 3’SL administration reduced atherosclerosis development as it would be a more clinically relevant therapeutic option. We determined absorption of orally administered 3’SL using different doses (3’SL, 30, 60, & 90 mg per mouse) into recipient wildtype mice (Fig. 7a). Oral administered 3’SL appeared in the blood stream within 5 min after the bolus, reaching a peak 20 min after ingestion and returned to baseline after 180 min. Oral delivery of 90 mg 3’SL resulted in peak concentrations of 9-12 µg/mL (Fig. 7a), which is in range of our effective IC50 dose tested *in vitro* (Fig. 1g). All tested concentrations were well tolerated and did not result in any undesirable gastrointestinal side effects. Therefore, we fed *Ldlr^-/-^* mice with a WTD two weeks prior to initiating a twice daily oral gavage of 90 mg/mouse 3’SL in 200 µl water, or 200 µl water as a control, for six weeks (Fig. 7b). 3’SL therapy did not alter food intake (Fig. 7c), body weight gain or organ weights of adipose tissues and liver at the time of harvest compared to control treated *Ldlr^-/-^* mice (Extended Data Fig. 6a-b). Plasma lipid levels and lipoprotein profile analysis showed a drop in plasma very low density lipoprotein (VLDL)-associated triglyceride and cholesterol levels in 3’SL treated mice (Fig. 7d-e and Extended Data Fig. 6c-d). However, no differences in the atherogenic LDL-cholesterol levels were observed between the treatment groups (Fig. 7e). Similarly, fasting blood glucose and hepatic lipid content were unaltered (Extended Data Fig. 6e-g).

**Figure 7:**
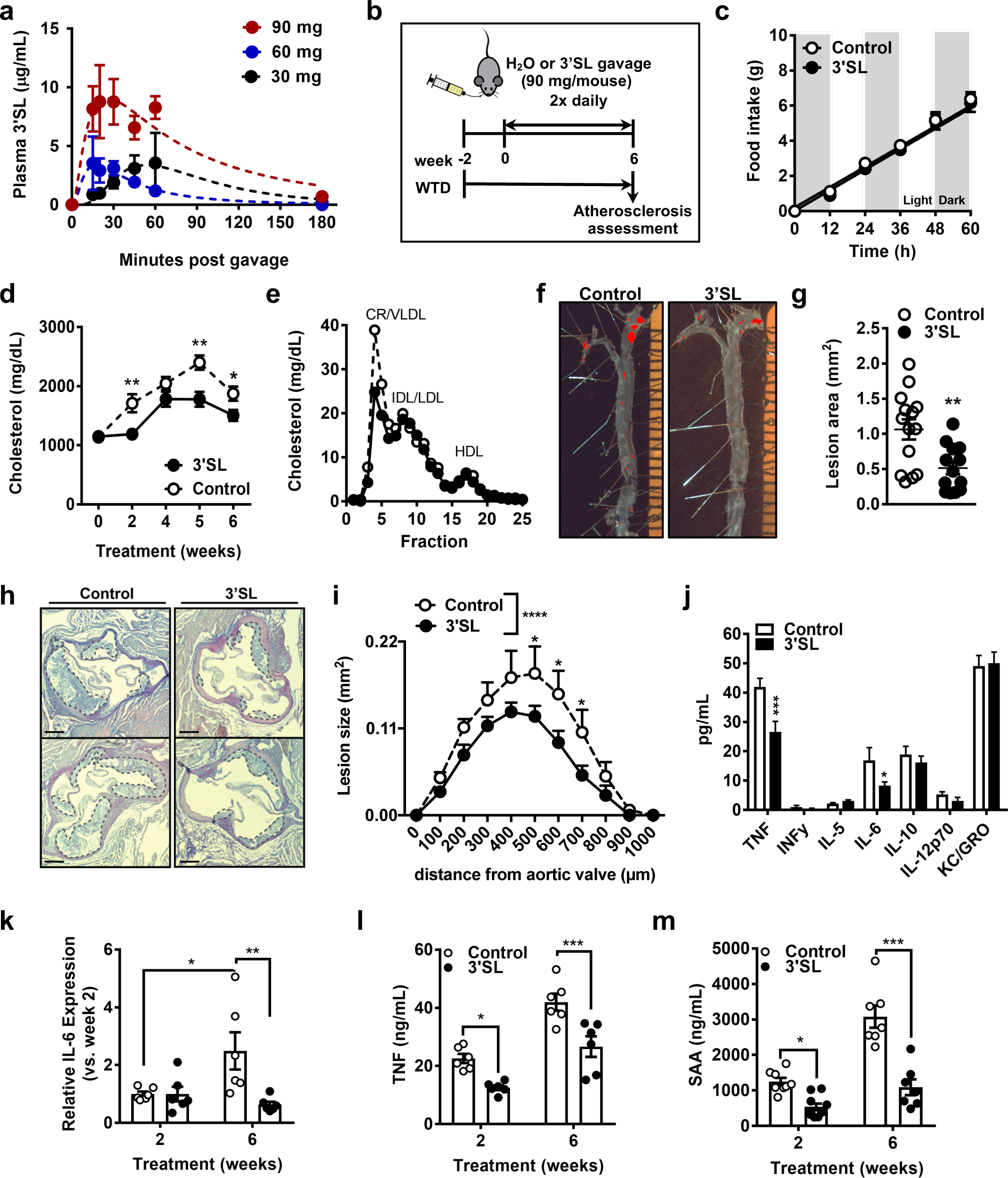
Six-week oral 3’SL treatment reduced atherosclerotic development and associated inflammation. **a**, Pharmacokinetics of 3’SL after oral injections (n = 3 per group/time point). **b**, Treatment regimen. Male *Ldlr-/-* were put on a Western type-diet (WTD) for 8 weeks. After two weeks, mice were treated twice daily with oral gavage of 90 mg 3’SL in water per mouse for 6 weeks. Water gavage served as a control. **c**, Food intake of *Ldlr-/-* mice on a Western type-diet measured 3 weeks into the treatment. **d**, Bi-weekly plasma cholesterol levels in 3’SL treated and control *Ldlr-/-* mice (n = 14-15). **e**, FPLC cholesterol lipoprotein profiles after 6 weeks of treatment (two pooled samples per group of 7-8). **f-g**, Representative Oil Red O staining (**g**) and *en face* quantification of atherosclerotic lesion size (**g**) in the aorta (n = 14-15). **h-i**, Representative H&E staining (**h**) and quantification of atherosclerotic lesion size (**i**) in the aortic sinus (scale bar = 100 µm, n = 14-15). **j**, Cytokine concentrations in the plasma after 6 weeks treatment (n = 6). **k**, Relative plasma IL-6 levels after 2 and 6 weeks of treatment (n = 6). **l**, Plasma TNF levels after 2 and 6 weeks of treatment (n = 7-8). **m**, Plasma SAA levels after 2 and 6 weeks of treatment (n = 6). (* p<0.05; ** p<0.01; *** p<0.001); Bar or dot plots represent mean ± SEM.

We next examined the impact of oral 3’SL administration on atherogenesis. Atherosclerosis lesion volume was assessed via *en face* aorta and aortic root analysis (Fig. 7f-i). Compared to control treated mice the six-week 3’SL administration resulted in a significant 51% reduction in *en face* atherosclerotic lesion area staining in the aorta (1.064 ± 0.1436 mm^2^ vs. 0.5129 ± 0.0898 mm^2^; *p* < 0.01) (Fig. 7f-g). In accordance, aortic root plaque analysis revealed that 3’SL treatment reduced lesion volume by 40% (1.0 ± 0.2 mm^3^ vs 0.6 ± 0.07 mm^3^; *p* < 0.0001) (Fig. 7h-i). Lesion analysis revealed no significant differences in necrotic core, collagen, macrophage, and smooth muscle cell content between the treatment groups (Extended Data Fig. 6g-j). These data imply that also oral 3’SL administration significantly attenuated development of WTD-induced atherosclerosis in *Ldlr^-/-^* mice.

### Oral 3’SL Administration in *Ldlr^-/-^* Mice Reduces Atherosclerosis-Associated Inflammation

In addition to lesion development, we were interested if atherosclerosis-associated inflammation was affected by oral 3’SL administration. Plasma cytokine analysis revealed that after six weeks of 3’SL treatment plasma tumor necrosis factor (TNF) and IL-6 levels were significantly reduced by 1.6- and 2-fold, respectively, compared to levels in *Ldlr^-/-^* treated controls (Fig. 7j). Two weeks into the treatment plasma IL-6 levels did not differ between 3’SL and control treated *Ldlr^-/-^* mice (Fig. 7k). After six weeks IL-6 levels significantly increased by 2.5-fold in control treated *Ldlr^-/-^*mice but showed a 1.5-fold decrease in 3’SL treated *Ldlr^-/-^* mice compared to plasma levels at two weeks (Fig. 7k). Oral 3’SL administration reduced plasma TNF by 1.8-fold already two weeks into the treatment and remained significantly lower throughout the treatment (Fig. 7l). In addition, we also measured serum amyloid A (SAA) in both 3’SL and control treated *Ldlr^-/-^* mice (Fig. 7m). SAA is an acute phase protein, and its plasma levels correlate with atherosclerotic development in mice and humans^63^. In mice, SAA serves as a proxy for the human atherogenic hsCRP, as CRP in mice is only a modest acute phase protein^64^. Oral 3’SL treatment was associated with significantly reduced plasma SAA levels after two and six weeks of treatment, by 2.3- and 2.8- fold respectively, compared to control treated *Ldlr^-/-^* mice (Fig. 7m). In conclusion, the findings suggest that oral 3’SL treatment reduces atherosclerosis development and attenuates atherosclerosis-associated inflammation in WTD fed *Ldlr^-/-^* mice.

## DISCUSSION

It is well established that activation of the innate immune response by TLR and inflammasome activation are crucial in the pathogenesis of atherosclerosis^42–44^. Currently, therapies aiming to reduce low-grade innate immune inflammation, in particular IL-1β targeting with monoclonal antibodies or decoy receptors, are being tested with success in clinical trials for their efficacy in reducing cardiovascular associated hospitalization and mortality^11, 12^. In this context we set out to test and identify the impact of specific HMOs, a group of natural compounds attributed with anti- and pro-inflammatory properties^35^, in macrophage inflammation. This study is the first to describe the impact of 3’SL on reducing low-grade chronic macrophage inflammation in the context of cardiovascular disease. Our data support that the anti-inflammatory properties of 3’SL are mediated via reduction of inflammatory gene expression, but equally by accelerating expression of genes in macrophages under control of LXR and SREBP that are important in the initiation of the inflammation resolution phase^58^. Based on the data, 3’SL treatment of macrophages appears to exert an atheroprotective role, by resolving inflammation and reducing the expression of chemoattractants and cytokines that are known to exponentially accelerate the development of atherosclerosis driven by hypercholesterolemia. Overall, our results support a concept that the HMO 3’SL can be used orally as a therapeutic in the context of adult diseases due to selective inhibitory effects on macrophage-driven low-grade chronic inflammation.

Several reports have described induction of TLR4 signaling by specific glycan structures, including HMOs^65^. The neutral HMOs LNFP II and LNnT affect peritoneal suppressor macrophages^66^, while other studies show that acidic HMOs have various immunomodulatory properties^37, 38^. In contrast, our study highlights the anti-inflammatory effects of 3’SL in LPS-stimulated macrophages. Our results suggest that 3’SL, and to as lesser extend 6’SL, are anti-inflammatory, yet the structurally related DSLNT is not. Both 3’SL and 6’SL are structurally almost identical with the only difference being the sialic acid moieties linkage to galactose which is an *α*2,3 and *α*2,6 linkage, respectively. Acidic HMOs such as 3’SL, 6’SL, and DSLNT share similar linkages, however DSLNT is sialylated by an *α*2,3 linkage to galactose and an *α*2,6 linkage to the internal *N*-acetyl glucosamine. Since each sialic acid contributes one negative charge of the HMO, the additional charge on DSLNT may interfere with its binding to a putative macrophage receptor. In addition, the molecular weight of DSLNT is nearly twice that of 3’SL or 6’SL (1,290.14 g/mol vs.

633.55 g/mol). The difference in size and charge of the structure may hinder the steric interactions of DSLNT and the macrophage receptor. Using genetic approaches, we also excluded that 3’SL anti-inflammatory properties are the result of simple engagement of the sialic acid with anti-inflammatory Siglecs expressed on macrophages^48^. The studies highlight that 3’SL and 6’SL have structure-specific anti-inflammatory properties that are poorly understood. However, simply carrying a sialic acid moiety on the HMO lactose backbone or complex HMO is not sufficient.

Previous reports have suggested 3’SL to inhibit NF-κB-p65 signaling in unstimulated Caco-2 cells, a cell model for enterocytes^67^. Also, a reduction in cytokine expression has been shown in cell types important for rheumatoid arthritis, such as T-cells, monocytes and osteoclasts^40^. At our effective dose we did not observe any impact of 3’SL administration on NF-κB p65 signaling. In addition to TRIF-signaling activation TLR4 stimulation induces a MyD88-independent signaling pathway that leads to the production of *IFN-β*^44^. However, 3’SL exposure did not affect *IFN-β* expression, *IFN-β* stimulated *Il-6* expression, nor did we observe changes in interferon-stimulated genes based on the transcriptome analysis. Instead, using an unbiased transcriptomic and epigenetic approach, we identified that the anti-inflammatory impact of 3’SL on TLR4 activated macrophages is surprisingly focused on a selected group of affected genes and enhancers. In contrast, over five thousand genes were differentially expressed in BMDMs in response to TLR4 activation independent of 3’SL co-incubation. This further supports the idea that 3’SL does not globally inhibit TLR4 activation and downstream NF-κB-p65 signaling. This concept is also corroborated by the observation that 3’SL attenuates TLR2-mediated macrophage activation. TLR2 activation is a well-established atherogenic pathway that primarily signals via the MYD88 pathway and does not require CD14 and MD2 co-receptors as is the case for TLR4 signaling^68^. Hence, our data also exclude that the anti-inflammatory effect of 3’SL is a consequence of altering signaling events mediated by CD14 or MD2 independently^69^.

Unexpectedly, we found that 3’SL was able to induce expression of a set of genes that is regulated by LXR and SREBP1, master regulators of cholesterol and fatty acid biosynthesis^60, 61, 70^. The modulation of lipid metabolism via these transcription factors is important for short term immune cell activation as well as long-term trained innate immunity^58, 71–74^. However, loss of function experiments indicated that LXRs and SREBPs were not required for the anti-inflammatory effects of 3’SL on LPS treated macrophages. Further studies will be required to determine the molecular mechanisms by which 3’SL activates LXR and SREBP and the functional consequences of these effects.

Studies of the effects of 3’SL on the epigenetic response to LPS revealed inhibition of H3K27ac and recruitment of the p300 histone acetyltransferase at a select set of enhancer elements associated with 3’SL-suppressed genes. These results imply a transcriptional mechanism by which 3’SL inhibits LPS-induced gene expression that involves direct or indirect effects on sequence-specific transcription factors and/or their coactivators. In-depth *de novo* motif analysis of 3’SL-repressed enhancers indicated that they were qualitatively different from LPS-induced enhancers that were not subject to 3’SL repression. Of particular interest, consensus binding sites for NFκB were present at less than 1% of 3’SL-sensitive enhancers, but direct ChIP-seq experiments indicated p65 occupancy at approximately 40% of these sites. In concert with the p300 binding profile, these findings suggest an indirect mode of NFκB binding at these enhancers that contributes to LPS activation but, in contrast to sites with consensus κB motifs, is sensitive to 3’SL repression. In addition to lack of consensus κB motifs, 3’SL-sensitive enhancers were distinguished from 3’SL-insenstive enhancers by the presence of several other transcription factor binding motifs. The most significantly enriched motif, for which a perfect consensus appears in 8% of 3’SL-sensitive enhancers, corresponds to a binding site for CTCF. CTCF is a highly conserved factor regulating gene transcription by spatially orienting *cis*-acting regulatory elements within topologically associating domains^75^. The strong enrichment of 3’SL-repressed genes for the CTCF motif raises the intriguing possibility that 3’SL alters LPS responses at specific genes by influencing CTCF-dependent interactions between enhancers and promoters. It will be of interest to investigate this possibility using chromatin conformation capture assays.

Several additional motifs were preferentially enriched in 3’SL-sensitive enhancers. The most frequently observed motif, found in 18% of 3’SL-sensitive enhancers, represents a binding site for IRF and STAT transcription factors. Enrichment for IRF motifs is particularly interesting as the 3’SL-repressed gene set is not associated with a canonical Type-I interferon signaling pattern of gene expression. BCL6 and RBPJ are also enriched in 3’SL inflammatory repressed genes and canonically serve as transcriptional repressors, that recruit additional TFs to form a co-repressor complex^76^. BCL6 encodes a BTB/POZ-zinc finger transcriptional repressor critical for the development and inflammatory potential of immune cells, including attenuation of NF-κB-p65 signaling in macrophages^76, 77^. BCL6 and the NF-κB cistromes intersect, within nucleosomal distance, at nearly half of BCL6-binding sites in stimulated macrophages to promote opposing epigenetic modifications of the local chromatin^78^. Recent studies in macrophages suggest that BCL-6 interacts with IκBζ and interferes its binding to the *IL-6* promoter in macrophages^79^. Further studies are needed to evaluate if 3’SL activates BCL6 recruitment directly or indirectly by activation of canonical ligand signaling pathways such as IL-4 and IL-21. However, BCL6 controls one-third of the TLR4 transcriptome^78^ and thus much more than the impact of 3’SL on TLR4 repression suggesting an additional level of regulation. RBP-J is a Notch signaling modulator and a regulator of pro-inflammatory macrophages polarization via activation of IRF8^80^. The latter might explain why we also see enrichment for the IRF motif, despite that fact that most Type-I IFN responsive genes are unaffected by 3’SL co-incubation (Extended Data Fig. 4b). It will be of interest to explore whether 3’SL can inhibit activation of the NOTCH:RBPJ TF complex and co-recruitment of co-activators and histone acetyltransferases to activate downstream inflammatory genes.

Importantly, we show that the anti-inflammatory effects of 3’SL *in vitro* translated in attenuated atherosclerosis lesion development *in vivo*. Anti-inflammatory effects of 3’SL *in vivo* were previously shown in mouse models of rheumatoid arthritis and atopic dermatitis^40, 81^. However, the authors administered 3’SL orally and thus it cannot be excluded that the anti-inflammatory phenotypes are due to likely drastic changes in the gut microbiome^34, 81^. As changes in the gut microbiome have been shown to alter the impact on atherogenesis^82, 83^, we intentionally sought to circumvent interactions of 3’SL with the microbiome and chose subcutaneous injections as our application route. Uptake of 3’SL into the circulation was efficient and the 6-week intervention did not result in any negative side effects, which was expected given the high tolerance and safety of the glycan^84–86^. After 8 weeks of WTD feeding whereby we treated for 6-weeks, *Ldlr^-/-^* mice have a reduction in lesion size and macrophage content within the lesions. This result is in line with *in vitro* reduced expression and secretion of pro-inflammatory cytokines and chemoattractants. In addition, we also observed that 3’SL administration reduces expression of endothelial chemokine and adhesion molecules the promote monocyte infiltration in cultured endothelial cells. It is very likely that both processes contribute to the attenuated atherogenesis and reduced plaque macrophage content. Based on previous reports we know that 3’SL does not directly compete with binding of monocytes to adhesion factors expressed by endothelium^37^. Hence, it will be important to probe in the future the impact of 3’SL administration on established atherosclerotic lesion regression and plaque rupture, which lead to relevant clinical events, in the future.

Finally, we also assessed the impact of oral 3’SL treatment on atherosclerosis development. Oral administration, which is clinically more relevant as a therapeutic strategy, reduced atherosclerosis even more strongly compared to a subcutaneous 3’SL intervention. A key difference between the two administration routes were triglyceride-rich lipoprotein levels, which were reduced by oral, but not by subcutaneous 3’SL treatment. While we observed no difference in LDL-cholesterol levels, the main atherogenic driver in this model, one cannot exclude that this reduction contributed to the anti-atherogenic effect of oral 3’SL treatment. However most triglyceride-rich lipoprotein lowering strategies fail to see an impact on atherosclerosis development in murine hypercholesterolemia models, unless they can also lower LDL-cholesterol levels^87^. Thus, it remains to be determined if this reduction contributes to the anti-atherogenic effect and whether the triglyceride-rich lipoprotein lowering effect is a result of reduced lipid absorption and VLDL production or accelerated hepatic clearance. The more profound anti-atherogenic effect induced by oral administration was also associated with a strong reduction of systemic inflammation. The observation that SAA went down a great deal upon oral 3’SL treatment is strong evidence for systemic decrease in inflammation as SAA is mainly made in the liver^64^. These favorable antiatherogenic discrepancy between the two administration routes could be a result of alterations in the microbiome in the gastrointestinal tract or due to a first pass effect of oral administered 3’SL via the liver, where the majority of the measured cytokines and triglyceride-rich lipoproteins are produced and cleared. Hence, upcoming studies will have to determine to what extend these differential and beneficial atherogenic outcomes associated with oral 3’SL administration are driven by microbiome alterations in the gastrointestinal tract versus direct anti-inflammatory effects on macrophages and endothelium.

The successful outcome of the CANTOS trial provides premise for reducing production of pro-inflammatory cytokines and thus the progression of the chronic inflammatory response as a secondary intervention strategy in patients with elevated hsCRP levels^11, 12^. With 3’SL being a natural compound, it could provide a viable alternative solution for treating chronic inflammation and promoting resolution of atherosclerosis and cardiovascular disease. Moreover, pro-resolving therapeutics are favorable over agents that directly block cytokine release, because it is expected that they are less likely to compromise the patient’s immune system. Until recently, clinical research studies have been limited by the insufficient availability of HMOs^88^. Due to advances in HMO isolation, purification and synthesis, new opportunities are surfacing for the large-scale HMO production, which guarantees the availability of HMOs for clinical trials and for commercial application^89–92^. Setting a precedent for 3’SL, the HMOs 2’FL and LNnT have already received GRAS (Generally Recognized As Safe) status by the United States Food and Drug Administration (FDA) for use in both infants and adults. In Europe, the same HMOs, have undergone novel food applications with the European Food Safety Authority (EFSA). Hence, our data warrant further analysis of 3’SL and its therapeutic potential in cardiovascular diseases and other chronic inflammatory disorders.

## ACKNOWLEDGMENTS

We would like to thank Jennifer Pattison and Karen Bowden for their assistance with aortic root sectioning. We would also like to thank UC San Diego Histology core facility for assistance with histology; the UC San Diego hematology core facility for hematology analysis; We thank Dr. Kristen Jepsen, for assistance with Illumina sequencing; and Leslie Van Ael for assistance with manuscript preparation. This work was supported by Foundation Leducq 16CVD01 (to P.L.S.M.G. and C.K.G.), NIH grant HD089067 (to P.L.S.M.G. and L.B.), DK113592 and HL140898 (to H.M.H.), HL 088093 (to J.L.W.), R35GM119850 (to N.E.L.), DK09118330 and DK063491 (C.K.G.), Metagenics Inc. (to L.B.), an Erwin-Schrödinger FWF Grant J4031-B21 (to A.R.P.), a Carlsberg Foundation Fellowship (to K.V.G.), and an American Heart Association (AHA) Predoctoral Fellowship 17PRE33410619 (to B.R.). L.B. is the UC San Diego Chair of Collaborative Human Milk Research endowed by the Family Larsson-Rosenquist Foundation, Switzerland.

## AUTHOR CONTRIBUTIONS

L.B. and P.L.S.M. conceived the project. A.R.P., N.J.S., C.A.A. H.M.H., N.E.L., J.L.W. C.K.G, L.B. and P.L.S.M. designed experiments. C.K.G., L.B. and P.L.S.M. supervised the project. A.R.P., N.J.S., C.A.A., B.R., J.L., A.S., K.V.G., A.W.T.C., N.E.L. and H.M.H. analyzed data. A.R.P., N.J.S., C.A.A., B.R, A.S., K.V.G., Y.W., A.Q., C.T., J.L., and L.M.B. performed experiments. A.R.P., N.J.S., C.A.A, C.K.G., L.B. and P.L.S.M.G. interpreted data and wrote the manuscript and the final versions were reviewed by all authors.

## METHODS

### Separation, purification and identification of HMO

Pooled HMO (pHMO) used for the experiments were isolated and prepared with a method described previously^93^. Breast milk collected at different times post-partum, was donated by healthy women whom gave birth at term. The milk from more than 50 women was pooled and centrifuged to remove the lipid layer. In addition, proteins were removed by precipitation from the aqueous phase with ice-cold ethanol. Ethanol was evaporated using a rotary evaporator and the remainder of the samples was lyophilized. Lactose and salts were removed by FPLC and samples were analyzed by HPLC, where HMO profiles containing less than 2% lactose were pooled and later used for experiments. Individual HMO used in experiments were either generously provided or purchased commercially (Extended Data Table 1).

Endotoxin (LPS) was removed from all pHMO and individual HMOs used for in vitro experiments by Detoxi-Gel Endotoxin Removing columns (Pierce Thermo Scientific, Rockford, IL, USA) according to the manufacturer instructions. The whole procedure was performed under sterile conditions with the use of pyrogen-free UltraPure distilled water, ethanol 200 proof and sterile filter tips to avoid LPS contamination. After collection of the flow-through, containing HMO, the purified samples were frozen immediately and lyophilized until completely dry. To minimize the amount of LPS, the HMO samples were processed twice on the endotoxin removing columns.

### Tissue culture

RAW 264.7 macrophages, HUVEC and THP-1 human monocytic cells were obtained from American Type Culture Collection (ATCC, Manassas, VA, USA). RAW264.7 cells were cultured in DMEM supplemented with 10% fetal bovine serum (FBS) and 0.1% Penicillin-Streptomycin (Pen-Strep) and THP-1 cells in RPMI supplemented with 10% FBS and 10 mM HEPES in a humidified incubator with a 5% CO2 at 37°C. Prior to experiments THP-1 cells were split into 12 well plates (1x10^6^ cells per well) and differentiated into macrophages by adding 600 nM phorbol 12-myristate 13-acetate (PMA) for 48 h. Differentiated macrophages attached to plate surface were used for experiments. HUVECs were grown in HUVEC, grown in 100-mm diameter dishes in EGM-2 medium (Lonza). BMDMs were isolated from femur and tibia from both hind legs of C57BL/6J mice and cultured in DMEM/F12 (supplemented with 10% FBS, 0.1% Pen-Strep) and 20 ng/mL MCSF to stimulate macrophage differentiation at 37°C, with 5% CO2. Medium with MCSF was refreshed every other day. On day 5 to 7 post isolation cells were seeded into 24-well plates (450,000 cells/well) and used on the following day for experiment. Bone marrow from *Siglece^-/-^ and Siglec1^-/-^* mice were a kind gift from Victor Nizet (UC San Diego). Peripheral blood monocytes (PBMCs) were isolated from small volumes of whole blood from healthy volunteers after informed consent, following a protocol for simple phlebotomy approved by the UCSD IRB/Human Research Protection Program, by Ficoll gradient and plated at 10^5^ cells per well in media without serum in a 96-well plate. Monocytes were allowed to adhere for 3 hours at 37°C before the supernatant was replaced with media with 10% FCS and incubated overnight at 37°C. LPS stock medium (10 ng/mL LPS in DMEM) was prepared and used for all cells activated with LPS at the day of the experiment. As a control for non-activated cells, PBS was used. HMOs diluted in H2O were added in the concentration needed. To avoid microbial contamination all prepared media were filter sterilized (0.22 μm filter units). When other stimuli were used to induce inflammatory responses, preparation of media and cell handling followed the procedure explained above.

### Multiplex Enzyme-Linked Immunosorbent Assay

BMDM were plated in 12-well plates (450,000 cells/well) as described above. Cells were co-incubated for 24 hours with 10 ng/mL LPS or PBS and 100 μg/mL 3’SL or PBS and supernatants were collected for measuring cytokine production of BMDM using the MSD Multi-spot Assay system Proinflammatory Panel 1 (V-Plex, K15048D, Mesa Scale Diagnostics, Rockville, MD, USA) or IL-6 or IL-1β (R&D systems) according to the manufacturer’s protocol. Cytokine levels in murine plasma were determined from EDTA-plasma from 4h fasted mice with the same V-Plex kit.

### Western Blot

Protein extracts were harvested with RIPA buffer (supplemented with proteinase inhibitor and PhosStop) from BMDMs seeded in 12-wells after indicated times of stimulation with PBS, LPS and 3’SL. The lysates were subjected to Western blot analyses (4-12% SDS-PAGE, 15 µg/lane) using phospho-p38 MAPK (Thr180/Tyr182, Cell signaling) and phospho-NFκB p65 (Cell signaling); as loading control antibody beta-Actin (Sigma Aldrich) was used.

### Cholesterol efflux assay

Cholesterol efflux from BMDMs was performed as described (Nakaya et al., 2011). Briefly, BMDMs were seeded in 48-well plates, labeled with 2 μCi/mL ^3^H-cholesterol. Twenty-four hours later, cells were washed and equilibrated overnight in DMEM/F12 with 0.2% bovine serum albumin (BSA fatty acid free). For the cholesterol efflux, the medium containing 25 μg/mL human HDL was added to cells. After 6-8 hours, aliquots of the medium were removed, and the ^3^H-cholesterol released was measured by liquid scintillation counting (LSC). The ^3^H-cholesterol present in the cells was determined by lysing the cells with 0.3N NaOH/0.1% SDS and measuring aliquots by LSC. The percentage cholesterol efflux was calculated by dividing the media-derived radioactivity by the sum of the radioactivity in the media and the cells.

### Mice

*Ldlr^-/-^* and C57BL/6 mice were purchased from The Jackson Laboratory and bred in our animal facility. *Lxrα/β* knock-out mice (*Lxr*^-/-^) were generated as described^94^. All animals were housed and bred in vivaria approved by the Association for Assessment and Accreditation of Laboratory Animal Care located in the School of Medicine, UCSD, following standards and procedures approved by the UCSD Institutional Animal Care and Use Committee. Mice were weaned at 4 weeks, maintained on a 12-hour light cycle, and fed ad libitum with water and standard rodent chow (PicoLab® Rodent Diet 20 5053) or a Western diet (TD.88137, Envigo Teklad) containing 42 % kcal from fat. Mice received 3’sialyllactose citrate in PBS (400 µg/injection) or PBS via subcutaneous injections twice daily for 6 weeks.

### RNA analysis

Total RNA was isolated from homogenized tissue and cells and purified using E.Z.N.A. Hp Total RNA kit (Omega Tek) or RNeasy Mini kit (Qiagen) according to the manufacturer’s instructions. The quality and quantity of the total RNA was monitored and measured with NanoDrop (NanoDrop Technologies, Inc. Wilmington, DE). For quantitative PCR analysis, 5-10 ng of cDNA was used for real-time PCR with gene-specific primers (Extended Data Table 2) and TBP as a house keeping gene on a BioRad CFX96 Real-time PCR system (Bio Rad).

### RNA sequencing library preparation

Transcriptomes of quiescent BMDM (PBS) were compared to BMDM incubated with either 3’SL or LPS alone and LPS and 3’SL together. RNA was extracted from cells and stranded RNA-Seq libraries were prepared from polyA enriched mRNA using the TruSeq Stranded mRNA library prep kit (Illumina). Library construction and sequencing was performed by the University of California San Diego (UCSD) Institute for Genomic Medicine. Libraries were single-end sequenced for 76 cycles on a HiSeq 4000 to a depth of 20-30 million reads.

### Chromatin immunoprecipitation with RNA-Seq (ChIP-Seq)

ChIP for H3K27ac was performed essentially as describe previously ^95^ with minor modifications. Briefly, 1x10E6 bone marrow derived macrophages (BMDMs) were fixed with 1% paraformaldehyde in PBS for 15 minutes at room temperature. Next, 2.625 M glycine was added to 125 mM to quench fixation and cells were collected by centrifugation with at 1,100 X G for 10 minutes at 4C. Cells were washed twice with cold PBS and collected by centrifugation at 1,100 X G for 10 minutes at 4C. Cell pellets were then snap frozen and stored at -80C. For ChIP reactions, cell pellets were resuspended in 500 µl swelling buffer (10 mM HEPES/KOH pH7.9, 85 mM KCl, 1 mM EDTA, 0.5% IGEPAL CA-630, 1x protease inhibitor cocktail, 1 mM PMSF) and incubated on ice for 5 minutes. Cells were spun down and lysed in 500 µl LB3 (10 mM Tris/HCl pH 7.5, 100 mM NaCl, 1 mM EDTA, 0.5mM EGTA, 0.1% deoxcycholate, 0.5% sarkosyl, 1x protease inhibitor cocktail (Sigma), 1 mM PMSF). Chromatin was sheared to an average DNA size of 100-400 bp by administering 6 pulses of 10 seconds duration at 12 W power output with 30 seconds pause on wet ice using a Misonix 3000 sonicator. Samples were collected and 10% Triton X-100 was added to 1% final concentration. From each sample, 1% was taken as input DNA. For chromatin immunoprecipitation, 30 µl of a 50:50 mixture of Dynabeads Protein A and Dynabeads Protein G coated with anti-H3K27ac (Active Motif) were incubated with slow rotation at 4°C overnight. For preparation of Dynabead-antibody complex, Dynabeads Protein A:G mixture and 4 μg specific antibody were incubated in 0.5% BSA/PBS for 1 hr at 4°C on rotator, then washed twice with 0.5% BSA/PBS and brought up to volume of 20 µl per IP with 0.1% BSA/PBS. After overnight incubation, the beads were collected using a magnet and washed three times each with wash buffer I (20 mM Tris/HCl pH 7.5, 150 mM NaCl, 1% Triton X-100, 0.1% SDS, 2 mM EDTA, 1x protease inhibitor cocktail, 1 mM PMSF) and wash buffer III (10 mM Tris/HCl pH 7.5, 250 mM LiCl, 1% Triton X-100, 0.7% Deoxycholate, 1 mM EDTA, 1x protease inhibitor cocktail, 1 mM PMSF). Beads were then washed twice with ice cold TE plus 0.2% Tween-20., then once with TE plus 50mM NaCl. Sequencing libraries were prepared for ChIP products while bound to the Dynabeads Protein A:G initially suspended in 25 µl 10 mM Tris/HCl pH 8.0 and 0.05% Tween-20. For LXR, SREBP, p65 and P300, ChIP-Seq with BMDMs was performed as described previously (Oishi et al., 2016, Sakai et al., 2020).

### ChIP-Seq Library preparation

ChIP libraries were prepared while bound to Dynabeads using NEBNext Ultra II DNA Library preparation kit as previously described^96^. Libraries were eluted and crosslinks reversed by adding to the 46.5 µl NEB reaction 20 µl water, 4 µl 10% SDS, 4.5 µl 5M NaCl, 3 µl 0.5 M EDTA, and 1 µl 20 mg/ml proteinase K, followed by incubation at 55°C for 1 hour and 65°C for 30 minutes to overnight in a thermal cycler. Dynabeads were removed from the library using a magnet and libraries cleaned by adding 2 µl SpeedBeads 3 EDAC in 61 µl 20% PEG 8000/1.5 M NaCl, mixing well, then incubating at room temperature for 10 minutes. SpeedBeads were collected on a magnet and washed two times with 150 µl 80% ethanol for 30 seconds. Beads were collected and ethanol removed following each wash. After the second ethanol wash, beads were air dried and DNA eluted in 13 µl 10 mM Tris/HCl pH 8.0 and 0.05% Tween-20. DNA was amplified by PCR for 14 cycles in a 25 µl reaction volume using NEBNext Ultra II PCR master mix and 0.5 µM each Solexa 1GA and Solexa 1GB primers. PCR amplified libraries were size selected 200-500 bp using gel extraction using 10% TBE acrylamide gels. Libraries were single-end sequenced using either a HiSeq 4000 or a NextSeq 500 to a depth of 10-20 million reads.

### Sequencing Data Analysis

FASTQ files from sequencing experiments were mapped to the mm10 assembly of the mouse genome. STAR with default parameters was used to map RNA-Seq experiments^97^. Bowtie2 with default parameters was used to map ChIP-seq experiments^98^. HOMER was used to convert aligned reads into “tag directories” for further analysis^62^. Each sequencing experiment was normalized to total of 107 uniquely mapped tags by adjusting the number of tags at each position in the genome to the correct fractional amount given the total tags mapped. Sequence experiments were visualized by preparing custom tracks for the UCSC genome browser ^99^ using pooled tag directories. Peaks from ChIP-Seq and ATAC-Seq were defined using findPeaks command with default criteria (4-fold enrichment over background tag count; poisson enrichment p-value < 0.0001 over background tag count) as specified in HOMER documentation. Where applicable, IDR^100^ was used to test for reproducibility between replicate experiments, and only peaks with IDR < 0.05 were used for downstream analysis. To quantify transcription factor binding and H3k27Ac at ATAC-defined accessible regions, peak files were merged with HOMER’s mergePeaks and annotated with raw tag counts using HOMER’s annotatePeaks using parameters -noadj, -size as indicated. Subsequently, DESeq2 ^101^ was used to identify the differential H3K27ac signal or chromatin accessibility with FC > 1.5. To identify motifs enriched in peak regions over the background, HOMER’s de novo motif analysis (findMotifsGenome.pl) including known default motifs and de novo motifs was used. The background sequences were either from random genome sequences or from peaks from the comparing condition indicated in the main text and in the figure legends. For RNA seq, each experiment was quantified using the “analyzeRepeats” script of HOMER. To generate a table of raw read counts, the parameters -count exons -condenseGenes -noadj were used. To generate a table of TPM values, the parameters -count exons -condenseGenes -tpm were used. The TPM values were further processed by log2(TPM+1). DE genes were identified using raw sequencing read counts by DESeq2 ^101^ analysis through the “getDifferentialExpression” HOMER command at p-adj (adjusted p value) < 0.05 and FC (fold change) > 1.5. Metascape was used for gene ontology analysis^102^.

### Ingenuity Pathways Analysis (IPA) upstream regulator and canonical pathway analysis

Upstream regulators and canonical pathways for LPS vs. LPS+3’SL, PBS vs. 3’SL, PBS vs. LPS+3’SL, and PBS vs. LPS were generated utilizing Ingenuity Pathway Analysis (IPA, QIAGEN Inc., https://www.qiagenbioinformatics.com/products/ingenuitypathway-analysis). Specifically, after Benjamini-Hochberg FDR correction, genes with adjusted p-values less than 0.05 and fold change greater than 1.5 were considered as differentially expressed genes (DEGs). For example, the list of 134 identified DEGs for LPS vs. LPS+3’SL generated a total of 180 predicted molecule types: ‘G-protein coupled receptor’, ‘cytokine’, and ‘other’, and dysregulated canonical pathways were deemed significant when reaching a |Z-score| >2. The same procedures were applied to the comparisons of PBS vs. 3’SL, PBS vs. LPS+3’SL, and PBS vs. LPS.

### Lipid and lipoprotein analysis

Lipid levels were analyzed in plasma and liver samples as previously described^87, 103^. Blood was drawn via the tail vein from mice fasted for 4-5 hr. Total plasma cholesterol and plasma triglyceride levels (Sekisui Diagnostics) and NEFA levels (WAKO Diagnostics) were determined using commercially available kits.

### Quantification of atherosclerotic lesions

Five hour fasted *Ldlr*^-/-^ mice were perfused with 10 ml of PBS following cardiac puncture. The heart and ascending aorta down to the iliac bifurcation were removed and incubated in sucrose buffer or formalin, respectively. The hearts were processed and stained by the atherosclerosis core morphology group at UC San Diego as described^47^. The isolated hearts were sectioned by cutting several 10 μm paraffin cross-sections starting with the first appearance of the first leaflet of the aortic valve until the last leaflet in 100 μm sections. Aortic root atherosclerosis was analyzed using modified Verhoeff-van Gieson elastic staining to enhance the contrast between the intima and surrounding tissues. At each 100 μm cross-section, the mean lesion size in each mouse was analyzed by computer-assisted morphometry (Image-Pro Plus 6.3, Media Cybernetics) by two investigators blinded to the study protocol. After removal of adventitial tissues, the aortas were incised longitudinally, pinned flat and stained for neutral lipids using Sudan IV. Images were acquired using a Leica MSV266 microscope with an attached Leica Ic80 HD camera.

### Histology of atherosclerotic lesions

Necrotic core formation was analyzed on modified Verhoeff-van Gieson elastic stained aortic root cross-sectional lesions (300 μm and 600 μm from beginning of aortic root) using ImageJ analysis software (NIH). Necrotic core area was measured as a percentage of total plaque area. Collagen content was quantified as percentage of total lesion area in aortic root cross-sectional lesions (300-400 μm from beginning of aortic root) stained with Picrosirius Red. Immunohistochemical analysis were performed on sections of paraformaldehyde-fixed and paraffin-embedded tissues. Aortic root cross-sectional lesions (300 μm from beginning of aortic root) were stained with anti-CD68 (Abcam, ab5694) and smooth muscle actin (Abcam, ab125212). The analysis of positively stained areas was done blinded by two investigators with the ImageJ analysis software.

### Statistics

Statistical analyses were performed using Prism software (version 7, GraphPad Software). Data were analyzed by unpaired 2-tailed Student’s *t* test or one-way ANOVA and presented as mean ± SEM. For experiments with more than two experimental groups, statistical significance was determined by ANOVA test for multiple comparisons. Dunnett’s post hoc tests were used for multiple comparisons to a control group and Bonferroni post hoc tests for multiple pair-wise comparisons. *P*-values are indicated by asterisk, where * *p*<0.05; ** *p*<0.01; *** *p*<0.001; and n.s. = not significant.

