## Supplemental file for "The Human Milk Oligosaccharide 3’Sialyllactose Promotes Inflammation Resolution and Reduces Atherosclerosis Development in Mice"

[1]Department of Medicine, [3]Department of Cellular and Molecular Medicine, [3]Department of Pediatrics, [4]Department of Bioengineering, [5]Novo Nordisk Foundation Center for Biosustainability, [6]Larsson-Rosenquist Foundation Mother-Milk-Infant Center of Research Excellence (MOMI CORE), [7]Rady Children's Hospital of San Diego, San Diego, CA, USA and [8]Glycobiology Research and Training Center, at University of California San Diego, La Jolla, CA, USA

***These authors contributed equally.**

**#To whom correspondence should be addressed:**

Lars Bode, Department of Pediatrics, University of California, San Diego, La Jolla, CA 92093-0687, Ph: 858/246-1874,

### EXTENDED DATA TABLES

**Extended Data Table 1:** Individually purchased human milk oligosaccharides.

| **Individual HMO** | **Company** |
|---|---|
| **3’sialyllactose sodium salt (3’SL)** | GeneChem |
| **6’sialyllactose (6’SL)** | Kyowa |
| **2’fucosyllactose (2’FL)** | Jennewein Biotechnologie |
| **Disialyllacto-N-tetraose (DSLNT)** | V-labs, INC |
| **Lacto-N-fucopentaose 1 (LNFP-1)** | V-labs, INC |

**Extended Data Table 2:** Primers used for qPCR analysis.

| Gene | Forward primer (5'-3') | Reverse primer (5'-3') |
|---|---|---|
| *CCL2*, human | TCTCAAACTGAAGCTCGCACTC | GCATTGATTGCATCTGGCTGAG |
| *Cxcl3,* murine | ATACTGAAGAGCGGCAAGTCC | AGACACCGTTGGGATGGATC |
| *Fdps,* murine | CTGGTGGTGCCAAGTGTGGAC | GAAGTGCTGGATGAAATTCTGC |
| *ICAM*, human | GGAGCACTCAAGGGGAGGTC | TGGCGGTTATAGAGGTACGTGC |
| *Il-10,* murine | TGAATTCCCTGGGTGAGAAG | TCACTCTTCACCTGCTCCACT |
| *IL-1β*, human | GTGGCAATGAGGATGACTTGTTC | TAGTGGTGGTCGGAGATTCGTA |
| *Il1-β*, murine | AAATACCTGTGGCCTTGGGC | CTTGGGATCCACACTCTCCAG |
| *IL-6*, human | AGCCACTCACCTCTTCAGAAC | GCCTCTTTGCTGCTTTCACAC |
| *Il-6*, murine | CCAGAGATACAAAGAAATGATGG | ACTCCAGAAGACCAGAGGAAAT |
| *IL-8*, human | AGCTGGCCGTGGCTCTCTTG | GGGTGGAAAGGTTTGGAGTATG |
| *Lrp8,* murine | AAGATTGAGAAGGCTGGGCTC | TACAAACGCTGGCTCAGCAAG |
| *Ptgs2,* murine | TGCACTATGGTTACAAAAGCTGG | CAGTCCGGGTACAGTCACAC |
| *Saa3, murine* | GTCCAGTTCATGAAAGAAGCTG | AGCATGGAAGTATTTGTCTGAG |
| *Stard4,* murine | CACAGCATCGAAGAAGATGAG | GTCATCCATAACTCCTTGAGC |
| *TBP*, human | CACGAACCACGGCACTGATT | TTTTCTTGCTGCTGCCAGTCTGGAC |
| *Tbp*, murine | GAAGCTGCGGTACAATTCCAG | CCCCTTGTACCCTTCACCAAT |
| *Tnf,* murine | ATTCGAGTGACAAGCCTGTAGC | GGTTGTCTTTGAGATCCATGCC |
| *VCAM*, human | GCGGGAGTATATGAATGTGAATC | AAGGAGGATGCAAAATAGAGCAC |

### EXTENDED DATA FIGURES AND LEGENDS

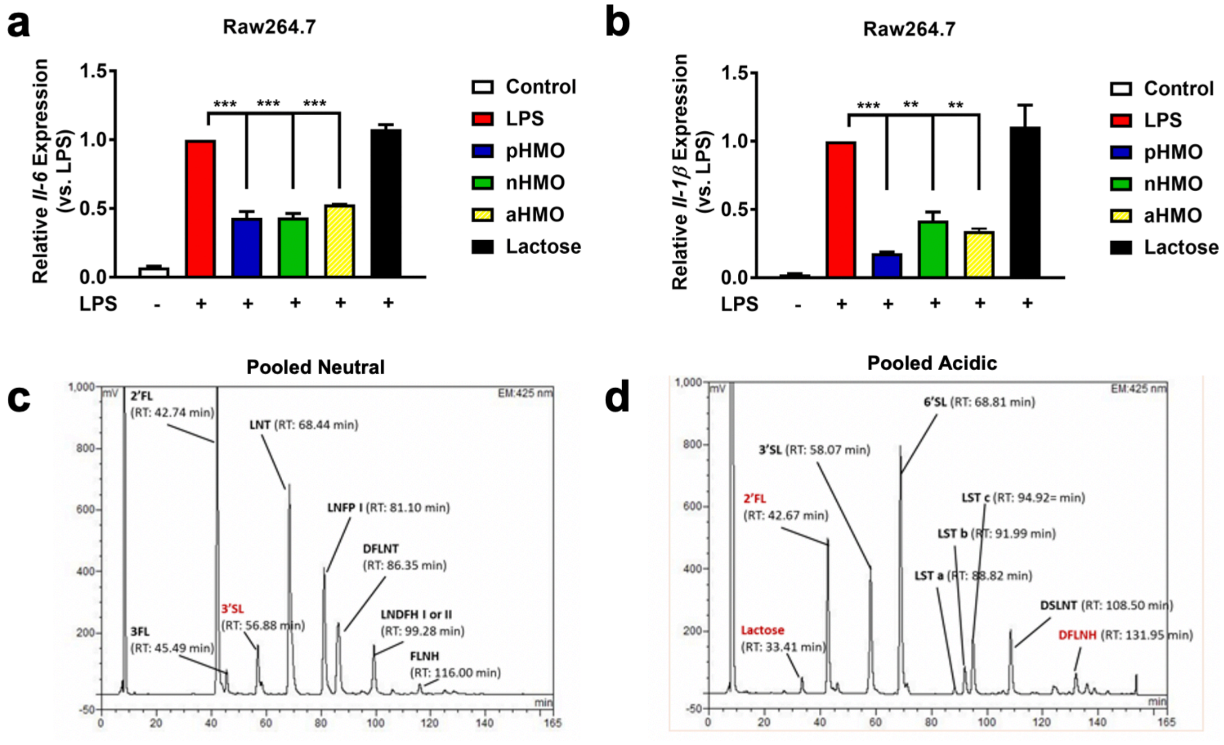


**Extended Data Figure 1. a-b**, Relative expression of *Il-6* (**a**) *and Il-1β* (**b**) in Raw264.7 cells when treated with LPS + pHMO, neutral (nHMO), acidic (aHMO), and lactose (all 500 μg/mL) (n = 2). **c-d**, Representative HPLC spectra of pooled (**c**) nHMO and (**d**) aHMO fractions. Statistical analysis with one-way ANOVA vs. LPS with Dunnett's post hoc test. (* $p<0.05$; ** $p<0.01$; *** $p<0.001$); Bar plots represent mean ± SEM.

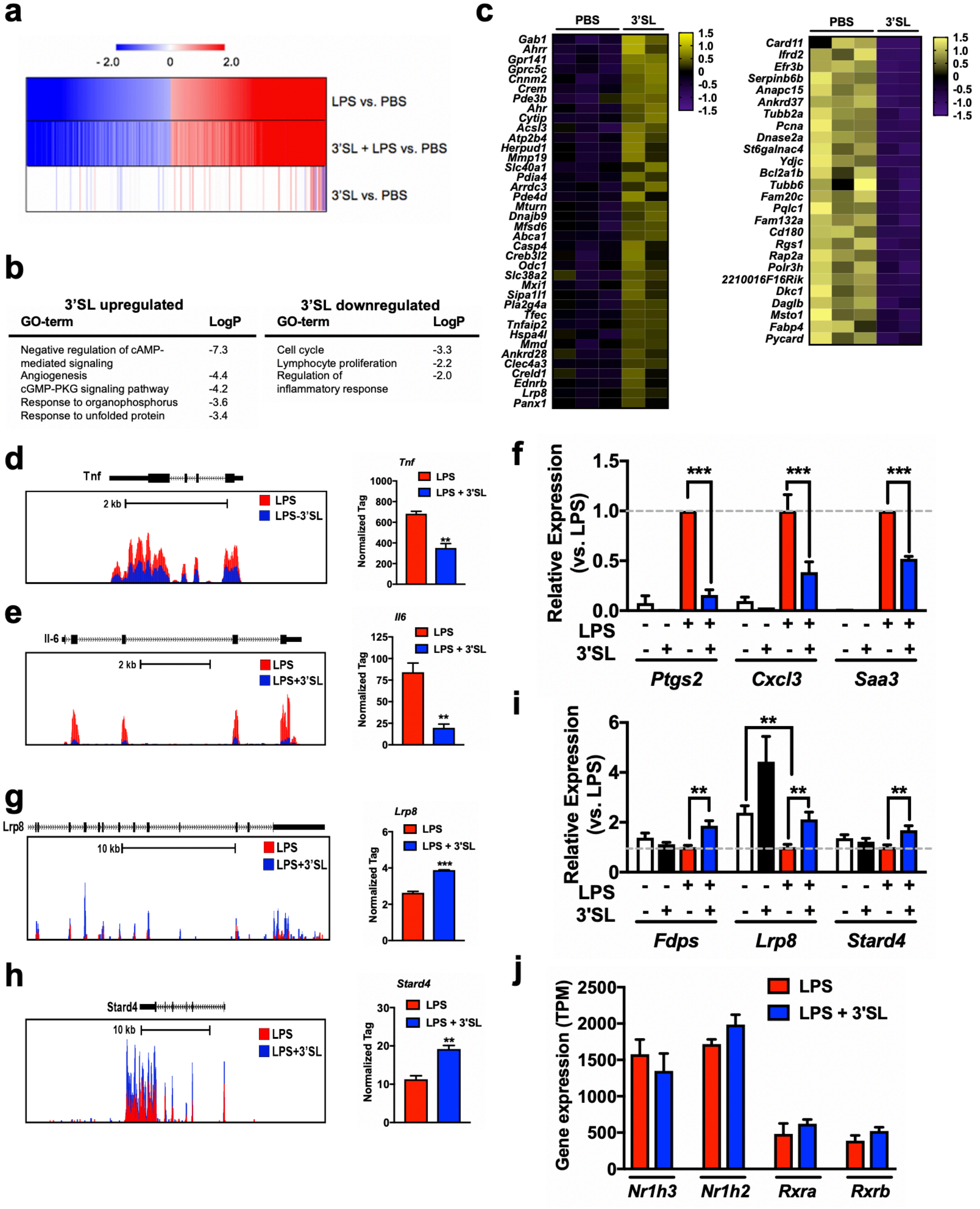


**Extended Data Figure 2. a**, Heat map representing z-normalized row expression of each gene for RNA-seq of all significantly up-regulated (red) or down-regulated (blue) genes in BMDMs at 6

h after 3'SL or LPS ± 3'SL stimulations in comparison with quiescent BMDMs (n = 3). **b**, Pathway analyses of 3'SL down-regulated and up-regulated genes in quiescent BMDMs. **c**, Heat maps representing fold-change expression of up-regulated (red) or down-regulated (blue) genes in quiescent BMDMs at 6 h after 3'SL stimulation (cut-off $p < 0.05$ and fold change, FC, >1.5) **d-e**, Normalized tag counts for *Tnf* and *Il-6* with normalized tag count averages (n=3). **f**, qPCR verification of *Ptgs2*, *Cxcl3* and *Saa3* in BMDMs (n ≥ 3). **g-h**, Normalized tag counts for *Lrp8* and *Stard4* with normalized tag count averages (n = 3). **i**, qPCR verification of *Fdps*, *Lrp8* and *Stard4* in BMDMs (n ≥ 3). **j**, RNA-seq analysis of LXR and RXR gene expression (n = 3). (** $p<0.01$; *** $p<0.001$); Bar plots represent mean ± SEM.

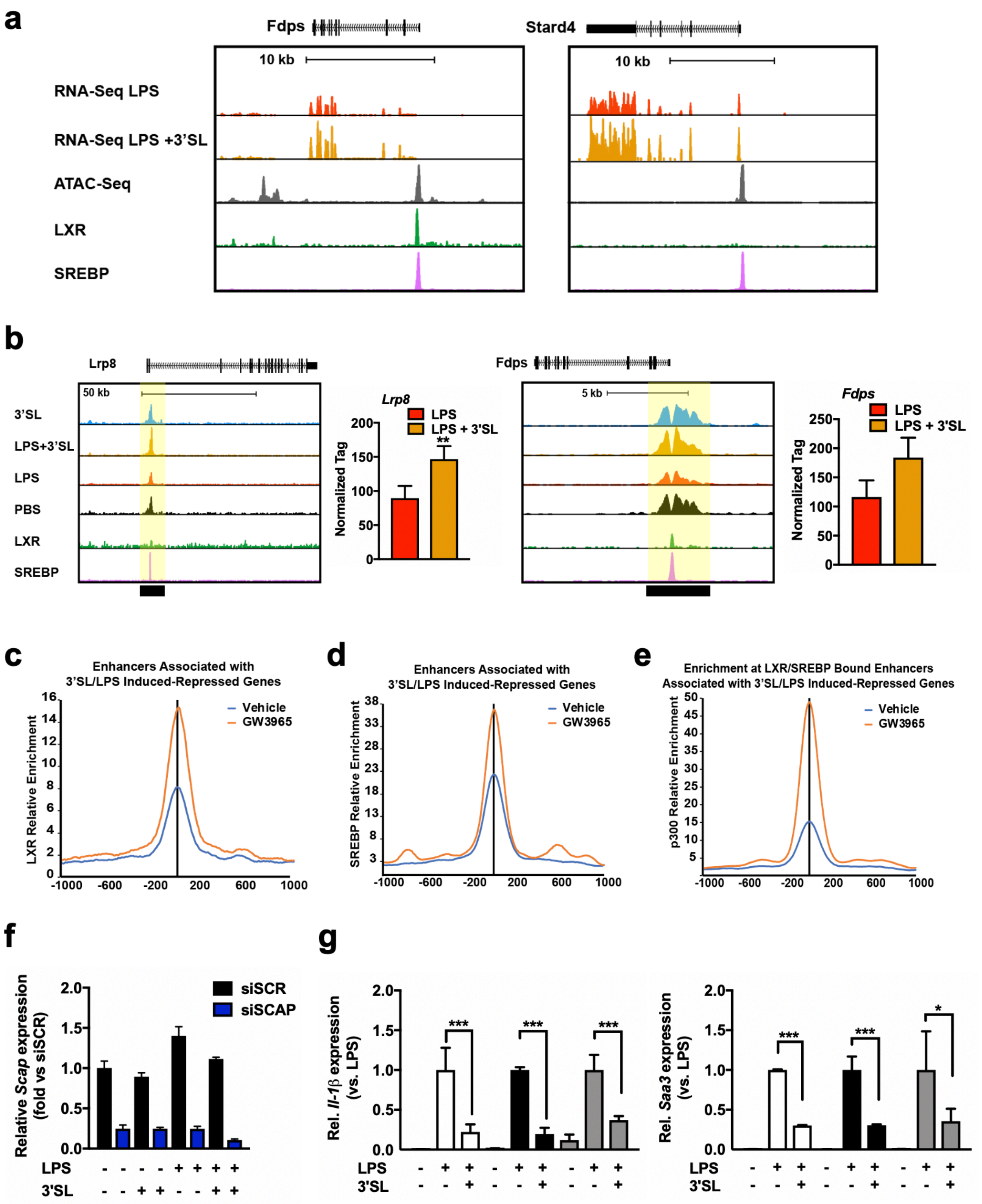


**Supplemental Figure 3 Related. a**, UCSC genome browser images illustrating normalized tag counts for *Fdps* and *Stard4* and illustrating ATAC-seq as well as mapped LXR and SREBP binding sites**.** **b**, UCSC genome browser images illustrating normalized tag counts for H3K27ac at 3'SL induced target loci together with mapped LXR and SREBP binding sites. **c-e**, Distribution of enhancers associated with 3'SL-LPS induced-repressed genes in the vicinity of genomic regions of vehicle and GW3965 treated macrophages co-bound by LXR (**c**) or SREBP (**d**) and p300 (**e**) enrichment at these sites. **f**, Relative *Scap* expression in BMDMs transfected with non-targeting control siRNA (siScr) or si*Scap* (n = 4). **g**, Relative expression of target genes in BMDMs transfected with non-targeting control siRNA (siScr) or si*Scap* or BMDMs isolated from *Lxrα/β* knock-out mice ($Lxr^{-/-}$) stimulated with PBS and LPS ± 3'SL. (n silencing experiments = 4, $Lxr^{-/-}$ BMDMs from 3 individual mice). (*p<0.05; ** p<0.01; *** p<0.001); Bar or dot plots represent mean ± SEM.

**a**

LPS 2h Induced Enhancers (vs PBS

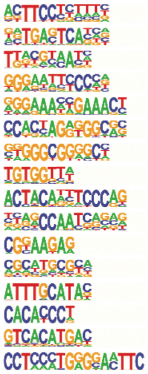

| Motif | p-value | %targ/ %bkgd | Best match |
|---|---|---|---|
| | 1e-2143 | 33.79/9.67 | PU.1(ETS)(0.963) |
| | 1e-1559 | 19.46/4.22 | Jun-AP1(bZIP)(0.992) |
| | 1e-382 | 17.74/8.88 | NFIL3(bZIP)(0.955) |
| | 1e-370 | 6.43/1.77 | NFkB-p65(RHD)(0.946) |
| | 1e-367 | 5.40/1.29 | STAT1/2(ISRE)(0.958) |
| | 1e-326 | 4.44/0.99 | CTCF(Zf)(0.948) |
| | 1e-319 | 16.35/8.48 | Sp1/2(Zf)(0.961) |
| | 1e-205 | 23.20/15.41 | RUNX1(Runt)(0.970) |
| | 1e-99 | 1.59/0.41 | Ronin(THAP)(0.916) |
| | 1e-98 | 6.78/3.88 | NFY(CCAAT)(0.934) |
| | 1e-93 | 11.25/7.41 | Gabpa(ETS)(0.763) |
| | 1e-71 | 3.10/1.46 | NRF1(NRF)(0.893) |
| | 1e-64 | 1.39/0.45 | Oct11(POU,Homeobox)(0.985) |
| | 1e-61 | 5.84/3.62 | KLF4(Zf)(0.959) |
| | 1e-54 | 2.79/1.41 | MITF(bHLH)(0.931) |
| | 1e-50 | 55.42/50.45 | RELB/NFkB-p65(RHD)(0.626) |

**b**

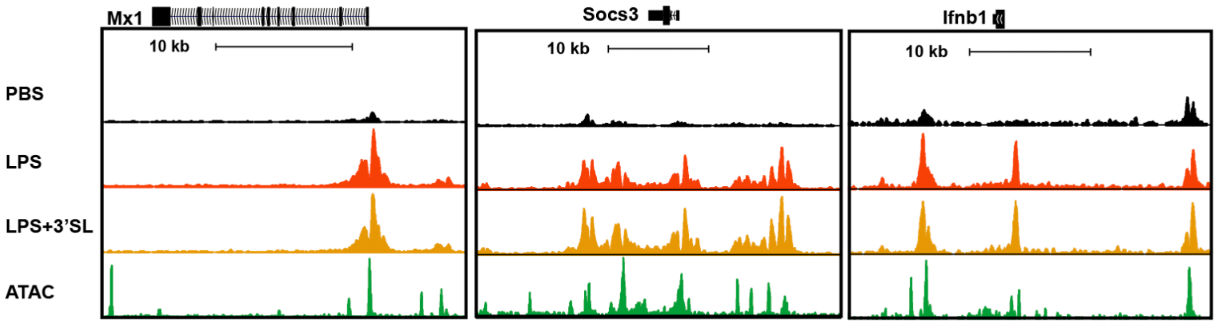


**Extended Data Figure 4. a**, *De novo* motif analysis of LPS upregulated enhancers (vs. LPS) using a GC-matched genomic background. **b**, UCSC genome browser images illustrating normalized tag counts for H3K27ac and illustrating ATAC-seq peaks at 3'SL induced target loci.

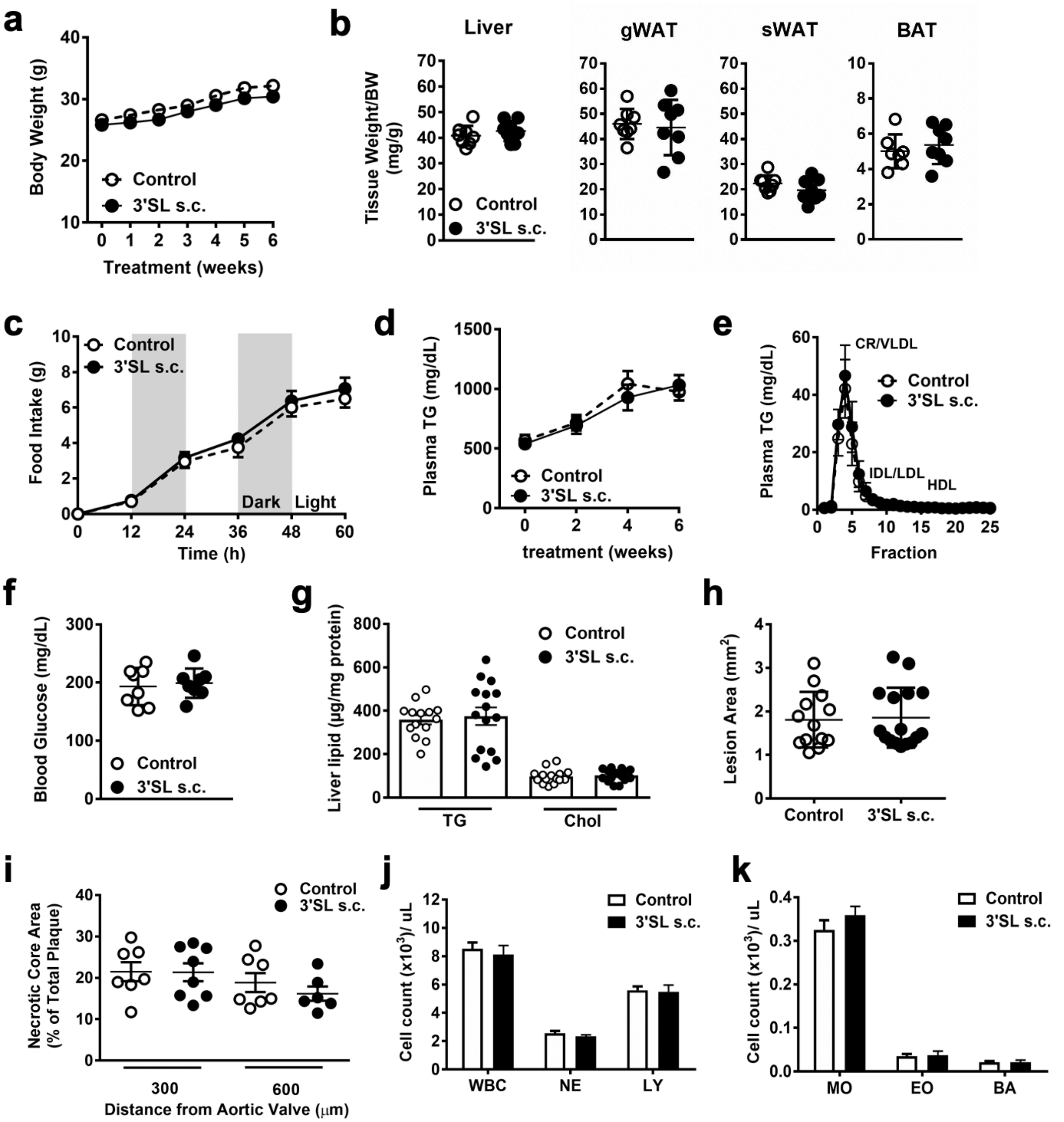


**Extended Data Figure 5. a**, Weekly body weight in subcutaneous (s.c.) 3'SL treated and control PBS treated *Ldlr*$^{-/-}$ mice (n = 8). **b**, Organ weights at harvest of Liver; gWAT, gonadal white adipose tissue; sWAT, subcutaneous WAT; BAT, brown adipose tissue (n = 8). **c**, Food intake over 60 hours measured every 12 hours. **d**, Bi-weekly plasma triglyceride (TG) levels in 3'SL treated and control *Ldlr*$^{-/-}$ mice (n = 14-15). **e**, FPLC TG lipoprotein profiles after 6 weeks of treatment (two pooled samples per group of 7-8). **f**, Plasma glucose was measured from full blood after 4 weeks of treatment (n = 8). **g**, Liver lipid levels after 6 weeks of treatment (n = 14-15). **h**, *En face* analysis of atherosclerosis and quantification of Sudan IV-positive area (n = 14-16). **i**, Quantification of necrotic core size (n = 7-8). **j-k**, White blood cell counts (WBC) in blood after 4 weeks of treatment. NE, neutrophils; LY, lymphocytes; MO, monocytes; EO, eosinophils; BA, basophils. All plasma parameters were measured after indicated treatment periods following a 5-hour food restriction. Bar or dot plots represent mean ± SEM.

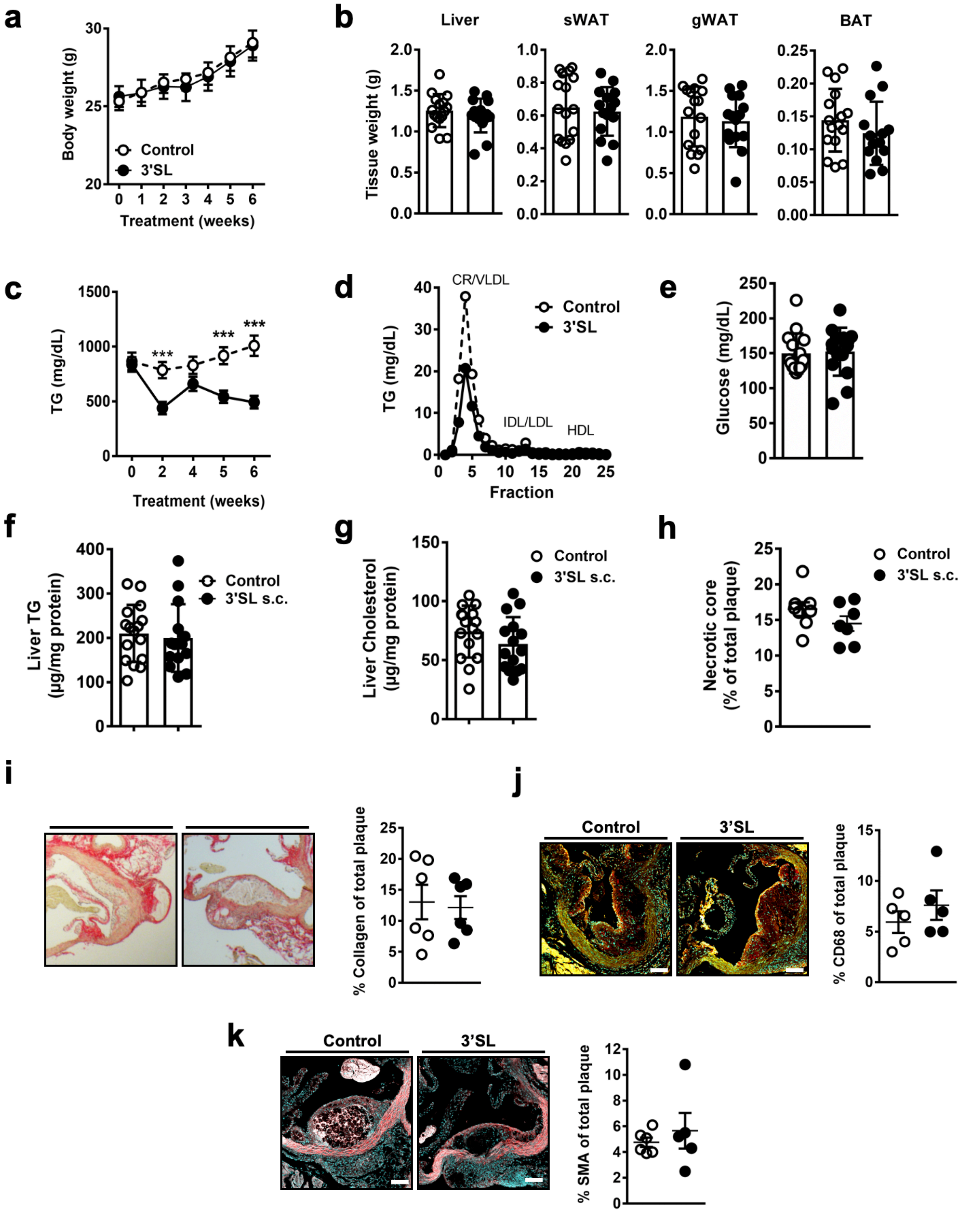


**Extended Data Figure 6. a**, Weekly body weight in oral 3'SL treated and control treated *Ldlr*$^{-/-}$ mice (n = 14). **b**, Organ weights at harvest of Liver; gWAT, gonadal white adipose tissue; sWAT, subcutaneous WAT; BAT, brown adipose tissue (n = 8). **c**, Bi-weekly plasma triglyceride (TG) levels in 3'SL treated and control *Ldlr*$^{-/-}$ mice (n = 14-15). **d**, FPLC TG lipoprotein profiles after 6 weeks of treatment (two pooled samples per group of 7-8). **e**, Plasma glucose was measured from full blood after 4 weeks of treatment (n = 8). **f-g**, Liver lipid levels after 6 weeks of treatment (n = 14-15). **h**, Quantification of necrotic core size (n = 7-8). **i**, Picrosirius red staining for collagen of equal sized atherosclerotic lesions and quantification of the positive stained area (n = 6). **j**, Atherosclerotic lesions stained with CD68 for macrophages and quantification of the positive stained area (n = 5). **k**, Smooth muscle actin (SMA) staining and quantification of the positive stained area (n = 5-6). All plasma parameters were measured after indicated treatment periods following a 5-hour food restriction. (*** $p<0.001$); Bar or dot plots represent mean ± SEM.
